# A pan-positive allosteric modulator that mediates sustainable GPCR activation

**DOI:** 10.1101/2025.03.24.645127

**Authors:** Qinxin Sun, Guodong He, Xinyu Xu, Fang Kong, Shuhao Zhang, Xiaoou Sun, Chuangye Yan, Xiangyu Liu

**Author notes:** Corresponding authors: Xinyu Xu and Xiangyu Liu. These authors contribute equally to the work: Qinxin Sun, Guodong He and Xinyu Xu. Department of Pharmaceutical Chemistry, University of California, San Francisco, San Francisco, CA 94143, USA.

## Abstract

Approximately 1/3 of all clinical drugs exert their therapeutic effects by modulating the activity of G protein-coupled receptors (GPCRs). Thus, there is a constant interest in finding novel ways to modulate GPCR activity. In this work, through a newly established Survival Pressure Selection (SPS) method for high-throughput screening of GPCR agonists, we discover that atazanavir functions as an agonist for GPR119. Further studies suggest that atazanavir is capable of activating a number of Family A GPCRs, including the β_1_ adrenergic receptor (β_1_AR), the β_2_ adrenergic receptor (β_2_AR) and the μ opoioid receptor (μOR). Cryo-EM structures reveal that atazanavir binds to an allosteric pocket near TM6/7 in both GPR119 and the β_1_AR. Pharmacological studies suggest that atazanavir mediates non-canonical signaling of Family A GPCRs. During this process, G protein and β-arrestin are spatially close, but β-arrestin forms a complex with the receptor and G protein instead of desensitizing G protein signaling, thus resulting in sustained G protein signaling. The work expands the signal transduction modes and pharmacological properties of allosteric modulators for GPCRs and provides a general starting point for developing therapeutics targeting this allosteric site.

## Main

G protein-coupled receptors (GPCRs) represent the largest family of cell surface receptors and the most successful class of drug targets^1^, owing to their crucial roles in a variety of physiological and pathological contexts. To date, approximately one-third of FDA-approved drugs exert their effects by modulating GPCR activity^2^. The development of innovative drug screening methods for GPCRs or the discovery of novel mechanisms for modulating GPCR function has profound implications for drug development and has attracted intense interest from both academia and industry.

Canonically, the activation of GPCRs triggers downstream G protein signaling, altering the cellular concentration of secondary messengers such as cAMP and Ca^2+^, thereby initiating different signaling cascades. Additionally, activated GPCRs can be phosphorylated by G protein-coupled receptor kinases (GKRs) to recruit β-arrestin1/2, typically leading to the desensitization of G protein signaling and receptor internalization^3,4^. Based on differences in receptor affinity for β-arrestin1/2, previous studies have classified GPCRs into Class A and Class B^5^. Class B GPCRs have a higher affinity for β-arrestin than Class A GPCRs. Certain Class B GPCRs, such as the parathyroid hormone type 1 receptor (PTHR)^6^, thyroid-stimulating hormone receptor (TSHR)^7^ and vasopressin type 2 receptor (V_2_R)^8^, have been reported to exhibit sustained G protein signaling over extended periods from internalized receptors. However, the therapeutic potential of the sustained intracellular activation remains poorly investigated, as this phenomenon appears to be an intrinsic property of Class B GPCRs with high affinity to β-arrestin. Achieving similar signaling behaviors for Class A GPCRs has been challenging, and it has not been previously reported that any ligand can mediate sustained intracellular activation of Class A GPCRs.

In this work, we established a new GPCR agonist screening method in *Saccharomyces cerevisiae* through protein fragment complementation techniques. Using this approach, we identified atazanavir, an FDA-approved drug for HIV treatment^9^, as a new positive allosteric modulator (PAM) for GPR119, an attractive therapeutic target for metabolic disorders such as type 2 diabetes^10,11^. Further pharmacological and structural investigations reveal that atazanavir can broadly activate various GPCRs, making it a pan-PAM. More significantly, by promoting the formation of GPCR - G protein - β-arrestin megacomplexes, atazanavir mediates sustained G protein signaling from the intracellular space, thus exhibiting the potential to expand the signaling mode for Class A GPCRs.

### Discovery of GPR119 agonists by a yeast survival pressure selection system

GPR119, also named as glucose-dependent insulinotropic receptor, is predominantly expressed in the pancreatic β-cells and the gastrointestinal tract, and it plays an important role in regulating glucose homeostasis^12^. GPR119’s significant physiological functions have attracted considerable attention from the drug discovery industry, leading to the development of many potent agonists for GPR119 by several pharmaceutical companies^13^. However, the clinical trial outcomes are not encouraging to date. The discovery of GPR119 agonists featuring novel scaffolds and different pharmacological properties may provide opportunities to develop better therapeutic candidates. Therefore, we tried to establish a high-throughput screening method for GPR119 agonists, initially based on the current GPCR signaling assays, like cAMP GloSensor or NanoBiT-based β-arrestin recruitment assays. However, these assays have their limitations, including high costs, challenging handling procedures, and a short signal detection time window.

To address these problems, we aimed to develop a new approach for GPCR agonist screening by converting the events of target GPCR activation to the downstream readouts of yeast growth. The assay is conducted in yeast *Saccharomyces cerevisiae*, which offers greater robustness for maintenance and manipulation compared to mammalian cells. The assay readout is the optical density (OD600) of the yeast culture, which remains relatively constant throughout the detection period. In brief, we employed a protein-fragment complementation strategy utilizing N-(5- phosphoribosyl)-anthranilate isomerase (Trp1p) as a reporter protein. Trp1p is an essential enzyme in yeast tryptophan biosynthesis and is indispensable for yeast proliferation in tryptophan-deficient environments. In our system, Trp1p is split into two components according to previous studies^14^. The N-terminal domain (NTD) of Trp1p is inserted into the α4-β6 loop of miniGs^15^ protein, while the C-terminal domain (CTD) of Trp1p is linked to the C-terminus of GPCR. Following agonist binding, the miniG protein is engaged by the activated receptor and enables the complementation of Trp1p, resulting in an increase in yeast growth rate in the selective medium lacking tryptophan **(Fig. 1a)**. The previously reported GPR119 agonist AR231453^16^ served as a positive control for system setup. After optimizing the linker for Trp1p NTD with miniGs and CTD with GPR119, along with the truncation site for the C terminus of GPR119, we obtained a construct that results in an obvious growth difference between the AR231453-treated condition and the untreated condition **(Extended Data Fig. 1a,c)**. Of note, so far, all the GPCR-G protein complexes share a similar interaction pattern between the receptors and the G proteins. As a result, this method is likely applicable to other GPCRs. Indeed, a similar strategy was applied to the β_2_AR and yielded a yeast strain that exhibits enhanced growth in response to β_2_AR agonists **(Extended Data Fig. 1b,d)**. The method was named the <u>S</u>urvival <u>P</u>ressure <u>S</u>election (SPS) method.

**Figure 1.**
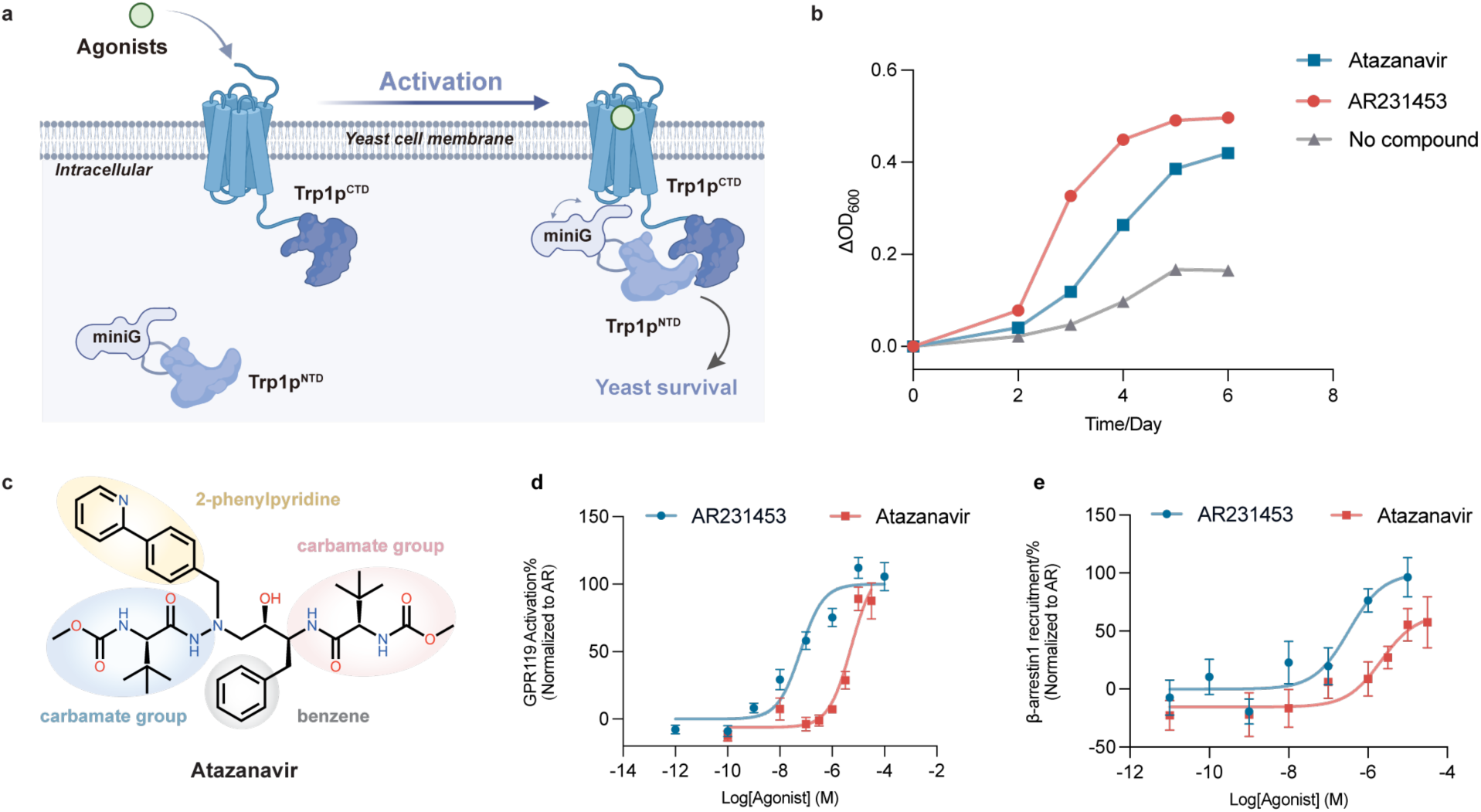
Identification of atazanavir as a GPR119 agonist by the SPS method. **a,** Design of the Survival Pressure Selection (SPS) system for GPR119, created with BioRender.com. **b,** The 6-day growth curves of GPR119-SPS yeasts in presence of AR231453, atazanavir or without agonist. **c,** The chemical structure of atazanavir. **d,** Atazanavir activates GPR119 in cAMP GloSensor assay. **e,** Atazanavir activates GPR119 in NanoBiT-based β-arrestin recruitment assay. Data are given as Mean ± S.E.M. of 6-12 independent replicates for panels d and e.

The SPS method was then employed to screen for GPR119 agonists within a library containing approximately 12,000 compounds. In brief, 50 nl of the compound and 40 μl of GPR119-SPS yeasts were transferred to 384-well plates and incubated in each well for 6 days. The OD600 of each condition was monitored using a microplate reader **(Extended Data Fig. 1g)**. The top 30 compounds that promoted yeast growth (ΔOD600 > 0.37) were selected for further characterization **(Extended Data Fig. 1e)**. An interesting compound named atazanavir was identified among the candidates **(Fig. 1b)**. Atazanavir is an FDA-approved drug for treating HIV infection^9^. It shows a novel scaffold compared to previously reported GPR119 agonists **(Fig. 1c)**. A cAMP-GloSensor assay and a NanoBiT-based G protein or β-arrestin recruitment assay were performed to evaluate the activation properties of atazanavir on GPR119. The compound exhibits agonist activity to GPR119, with EC_50_ of about 3 μM in all three assays **(Fig. 1d,e and Extended Data Fig. 1f)**.

### Structure determination of GPR119 bound with AR231453 and atazanavir

To further understand the molecular mechanism by which atazanavir activates GPR119, we tried to solve the cryo-EM structure of active state GPR119 bound with atazanavir. The synthetic agonist AR231453 was also included in the structural studies. Inspired by previous studies^17^, a GPR119-miniGsiN construct was generated by fusing miniGsiN to the C-terminus of GPR119. The purified GPR119-miniGsiN protein was then mixed with Gβγ, Nb35 and scFv16 to form a more stable complex. Finally, we solved the structures of GPR119-AR231453 and GPR119-atazanavir complexes with 2.98 Å and 3.33 Å resolutions, respectively **(Extended Data Fig. 2,3 and Extended Data Table 1)**. The clear electron densities enabled modeling of the ligands and the majority of receptors except for ECLs 1 and 3 (residues 66-72 for the GPR119-AR232453 structure, residues 68-74 and residues 250-260 for the GPR119-atazanavir structure) and ICL3 (residues 209-220 for the GPR119-AR232453 structure and residues 210-217 for the GPR119- atazanavir structure). AR231453 binds to the orthosteric pocket, while atazanavir binds to an allosteric pocket near TM6/7 **(Fig. 2a)**, as will be discussed in detail later.

**Figure 2.**
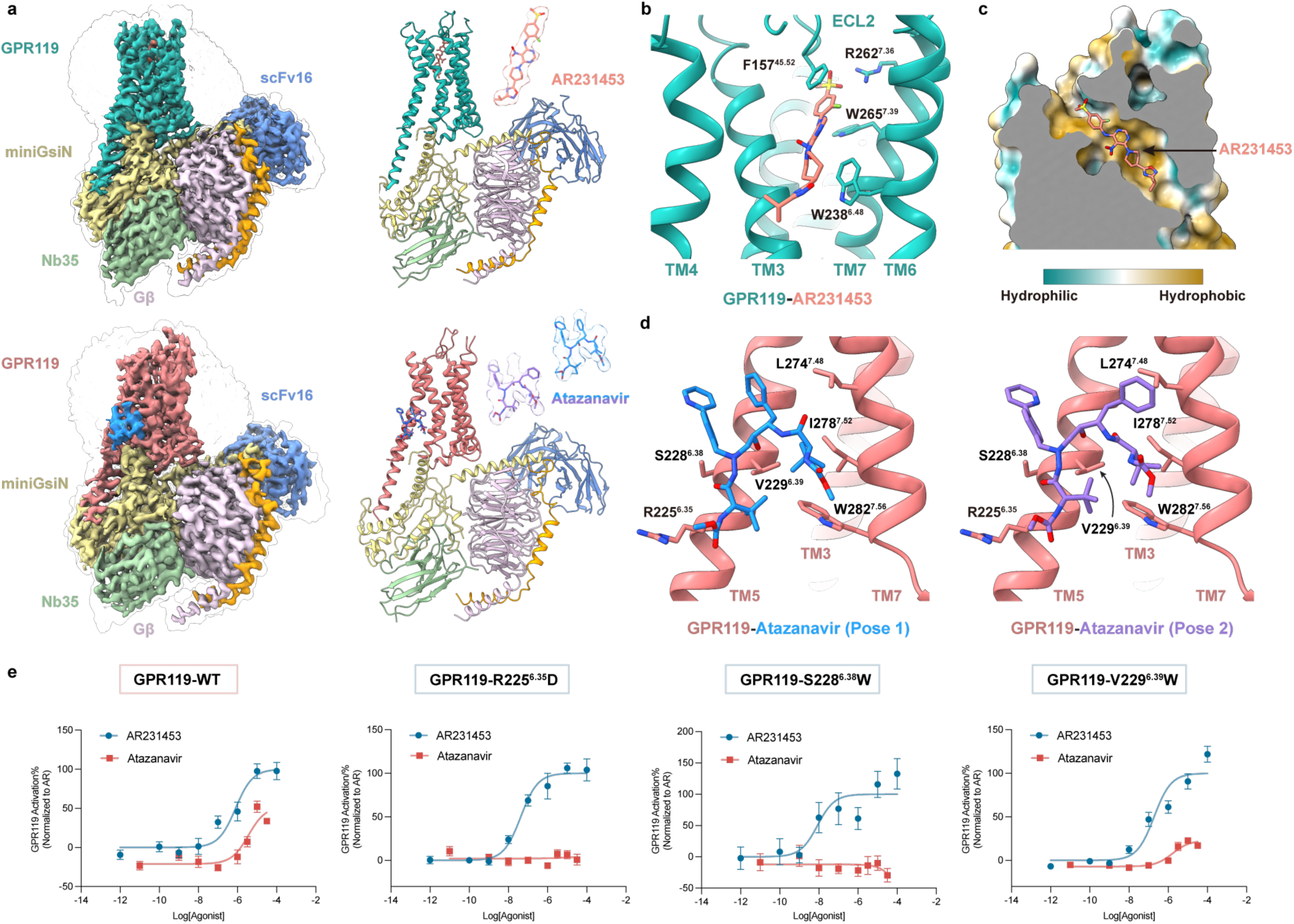
Cryo-EM structures of AR231453 and atazanavir-bound GPR119-Gs complexes reveal molecular details of ligands binding pocket. **a,** Overall structures of GPR119-miniGsiN-Gβγ-Nb35-scFv16 bound with AR231453 (top) or atazanavir (bottom). From left to right: electron density map, coordinate model and a zoom-in view of the electron densities for the ligands. AR231453-bound GPR119, dark green; atazanavir-bound GPR119, red; Gαs, yellow; Gβ, violet; Gγ, orange; scFv16, cornflower blue; Nb35, light green; AR231453, salmon; atazanavir (pose 1), blue; atazanavir (pose 2), purple. **b,** The ‘vertical’ binding pose and key interactions of AR231453 in the orthosteric pocket of GPR119. **c,** The hydrophobic (yellow) ligand binding pocket of AR231453. **d,** Atazanavir resides in a shallow pocket between TM6 and TM7 of GPR119. For both binding pose 1 and 2, the TM6 side of the binding pocket is formed by residues R225^6.35^, S228^6.38^ and V229^6.39^. **e,** The R225^6.35^D, S228^6.38^W and V229^6.39^W mutations impair atazanavir’s agonist activity for GPR119 but have little effect on AR231453’s agonist activity. Data are given as Mean ± S.E.M. of 10-12 independent replicates.

### The binding of orthosteric agonist AR231453

As the first potent and orally available GPR119 agonist discovered by Arena Pharmaceuticals, Inc., the basic pharmacological properties of AR231453 have been extensively studied^16^. Despite variations in sample preparation, our GPR119-AR231453 structure is very similar to a recently reported GPR119-AR231453 structure^18^. The structural information is consistent with the previous mutagenesis studies and pharmacological analysis. AR231453 binds ‘vertically’ at the canonical orthosteric binding pocket of Family A GPCRs **(Fig. 2a,b)**. The isopropyl-oxadiazole moiety penetrates deep into the binding core, and the 2-fluoro-4- methanesulfonyl-phenyl tail points toward the extracellular loops 2 (ECL2). The interactions between AR231453 and GPR119 are primarily hydrophobic **(Fig. 2c)**. The phenol ring of 2-fluoro-4-methanesulfonyl-phenyl moiety forms π–π interactions with the surrounding aromatic residues W265^7.39^ and F157^45.52^ (superscripts indicate Ballesteros–Weinstein numbering^19^). The W238^6.48^ also forms hydrophobic interactions with the isopropyl-oxadiazole moiety of the compound. While the oxygen atom of the SO_2_ tail forms a salt bridge with R262^7.36^ in TM7 **(Fig. 2b)**. These key interactions identified in the cryo-EM structure agree with prior studies that report W265^7.39^A and W238^6.48^A mutations diminish AR231453 activation, and R262^7.36^A and F157^45.52^A mutations cause a 6- to 79-fold decrease in AR231453 affinity^20^.

### Atazanavir is a positive allosteric modulator for GPR119

We also obtained a high-resolution structure of the GPR119-atazanavir complex (3.33 Å) and were surprised to find an absence of atazanavir-shaped density in the orthosteric pocket **(Extended Data Fig. 4a).** A narrow and continuous electron density was observed in the middle of transmembrane helices. The observed density is most likely contributed by the endogenous lysophospholipids, as reported in a recent study^21^. A unique density with the shape of atazanavir was observed near the membrane-facing surface of the cytoplasmic end of TM6/7. Model building suggested this density might represent an average of two possible binding poses of atazanavir (pose 1 and 2), with the benzene group adopting different conformations **(Fig. 2a)**.

To confirm the observed density was contributed by atazanavir, we performed mutagenesis studies on the residues near the atazanavir-shaped density. The residues on TM7 contribute to forming the atazanavir binding pocket, but their side chains do not directly orient towards atazanavir. We then mutated residues on TM6, including R225^6.35^, S228^6.38^ and V229^6.39^ **(Fig. 2d)**. The S228^6.38^W and R225^6.35^D mutations diminish the potency of atazanavir with little effect on the orthosteric agonist AR231453. The V229^6.39^W also decreases the maximal response of GPR119- mediated cAMP accumulation for atazanavir, likely due to steric hindrance **(Fig. 2e)**. Of note, these mutations have comparable expression levels as the wild-type receptor **(Extended Data Fig. 4d)**. Interestingly, we failed to identify specific interactions between atazanavir and the binding pocket residues, except for a potential hydrogen bond between R225^6.35^ and the methyl carbamate group of atazanavir. Atazanavir appears to reside in this shallow pocket between TM6 and TM7, as the pocket and compound geometries align remarkably well. It should be mentioned that atazanavir is sandwiched between TM6/7 and the phospholipids of the membrane bilayer. The atazanavir binding site includes a portion of hydrophilic surface **(Extended Data Fig. 4c)**. The direct exposure of this hydrophilic area to the hydrophobic membrane environment would be energetically unfavorable, while atazanavir binding shields this hydrophilic surface, thereby stabilizing the interaction. This may explain the μM range potency of atazanavir on GPR119 despite the lack of specific interactions between the compound and the receptor.

We then examined whether atazanavir could allosterically modulate the activity of AR231453. In a cAMP-GloSensor assay, the allosteric effect of atazanavir results in a 3-fold improvement of EC_50_ and a 2-fold increase in E_max_ for AR231453 at GPR119 **(Extended Data Fig. 4e)**. Similar allosteric effects of atazanavir are also observed for the endogenous agonist OEA **(Extended Data Fig. 4f)**. The enhancement in potency supports that atazanavir functions as a positive allosteric modulator for GPR119.

### Atazanavir serves as a pan-PAM for Family A GPCRs

The outward displacement of TM6 is a hallmark of GPCR activation^22^. Atazanavir recognizes and binds to the outwardly displaced conformation of TM6 **(Extended Data Fig. 4b)**, thereby increasing the population of active state receptors. Previous studies reported two allosteric modulators for GPR88^23^ and glucagon-like peptide-1 receptor (GLP-1R)^24^ that also bound to the intracellular surface of TM6 **(Extended Data Fig. 5a,b)**. This suggests that this position may represent a general allosteric modulation site for GPCRs. The sequence alignment of Family A GPCRs reveals that the residues surrounding the atazanavir’s binding site are relatively conserved, suggesting the possibility of atazanavir serving as a versatile agonist for Family A GPCRs **(Fig. 3a)**.

**Figure 3.**
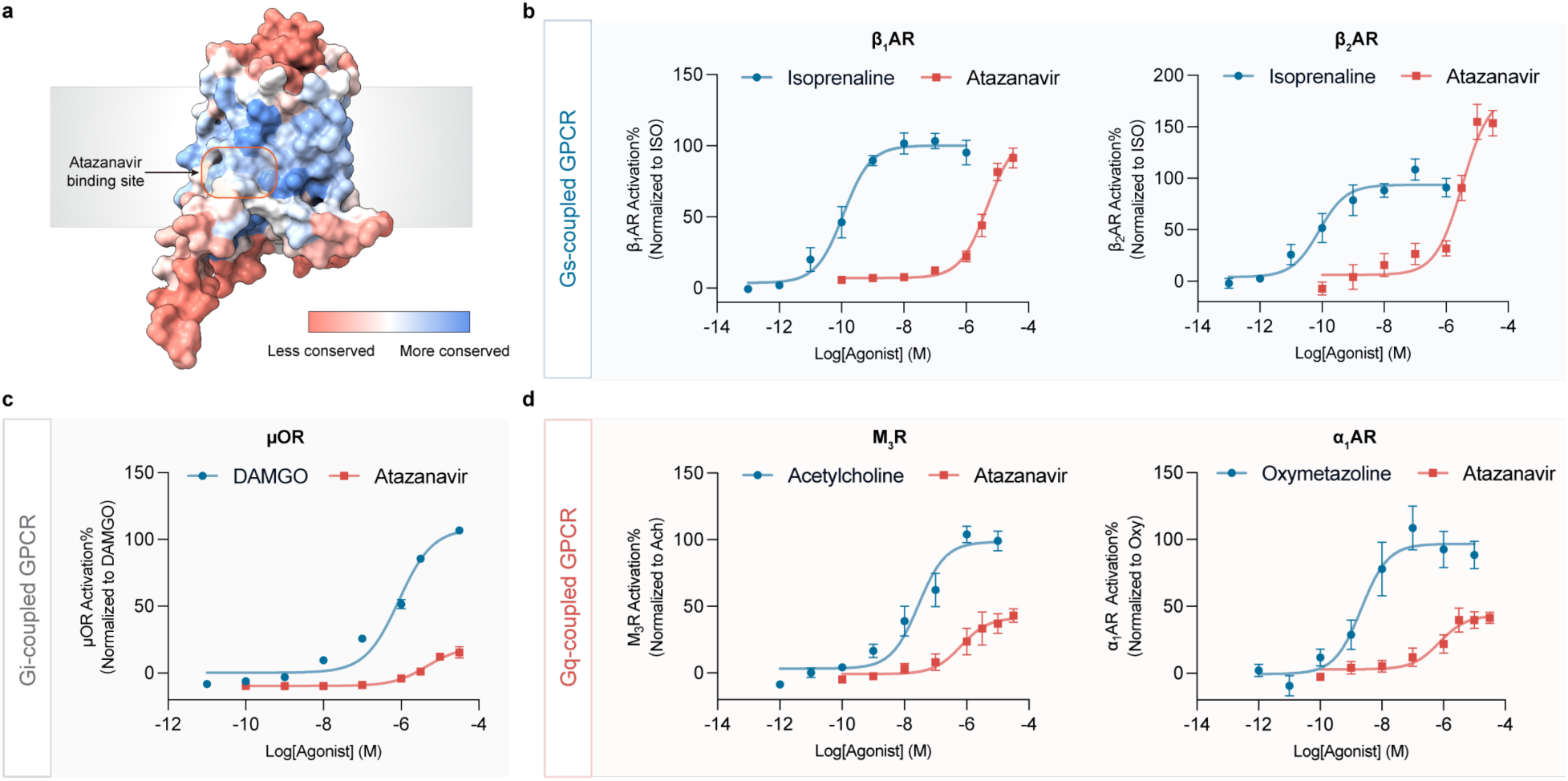
Atazanavir is a pan-PAM for Family A GPCRs. **a,** The atazanavir’s binding site is relatively conserved in Family A GPCRs. The sequence conservation analysis is done by GPCRdb (https://gpcrdb.org). **b,** Atazanavir activates Gs-coupled GPCRs (β_1_AR and β_2_AR) in cAMP GloSensor assay. **c,** Atazanavir activates Gi-coupled GPCR (μOR) in NanoBiT-based G protein recruitment assay. **d,** Atazanavir activates Gq-coupled GPCRs (M_3_R and α_1_AR) in cAMP GloSensor assay in the presence of a chimeric Gsq protein. Data are given as Mean ± S.E.M. of 6-9 independent replicates for panels b-d.

To assess the potential of atazanavir to modulate the function of other Family A GPCRs, we selected several representative receptors that couple to different types of G protein. Out of these 10 Family A GPCRs we tested, atazanavir exhibits agonist activity towards 8 of them, while dopamine D1 receptor (D_1_R) and neurokinin 2 receptor (NK2R) do not respond to the compound **(Extended Data Fig. 5c)**. We chose 5 receptors (β_1_AR, β_2_AR, μOR, M_3_R, ⍺_1_AR) for in-depth analysis. For Gs protein-coupled receptors, we utilized the cAMP GloSensor assay to verify atazanavir-induced receptor activation. The results demonstrate that atazanavir can activate both the β_1_AR and β_2_AR with EC_50_ values in the μM range, achieving efficacy that is comparable to or exceeds that of the full agonist isoprenaline **(Fig. 3b)**. The NanoBiT-based miniGs recruitment assay corroborate this finding, confirming the activation of the β_1_AR and β_2_AR by atazanavir **(Extended Data Fig. 5d)**. In the case of Gi protein-coupled μOR, the NanoBiT-based miniGi recruitment assay reveals that atazanavir promotes the interaction between μOR and miniGi, despite the lower efficacy compared to the orthosteric agonist DAMGO **(Fig. 3c)**. For Gq protein-coupled receptors, we employed the cAMP GloSensor assay using a Gsq chimeric protein, as well as the NanoBiT-based miniGsq recruitment assay. These assays indicate that atazanavir functions as a partial agonist for two representative Gq-coupled receptors, the M3 muscarinic acetylcholine receptor (M_3_R) and the α1 adrenergic receptor (α_1_AR), although its potency is low **(Fig. 3d and Extended Data Fig. 5d)**. Furthermore, for all the aforementioned targets, atazanavir exhibits cooperative effects when combined with their orthosteric agonists, resulting in enhanced EC_50_ and E_max_ values **(Extended Data Fig. 5e)**. However, as mentioned earlier, it is important to note that not all tested receptors can be activated by atazanavir. For example, D_1_R is not activated by atazanavir **(Extended Data Fig. 5c)**, likely due to the absence of suitable binding sites for atazanavir in this receptor.

Taken together, our results suggest that atazanavir demonstrates positive allosteric modulator activity on different GPCRs and can thus be termed a pan-PAM for these Family A GPCRs, making it a valuable tool for investigating the function of various receptors. The reasons for its versatility are multifaceted. Firstly, atazanavir contains several rotatable single bonds, enabling it to adopt multiple conformations suitable for different binding pockets. Secondly, the residues in atazanavir’s potential binding pocket near TM6/7 are relatively conserved among Family A GPCRs. More importantly, the conformational change of TM6 is a general feature of GPCR activation, and atazanavir binds to this outward-movement state, thereby promoting receptor activation.

### Structure determination of atazanavir binding to the β_1_AR

To deepen our understanding of how atazanavir recognizes and activates different receptors, we attempted to determine the structures of atazanavir bound to other GPCR-G protein complexes. The human β_1_AR was selected for structural investigation because of its good behavior and important physiological role. The wild-type human β_1_AR was purified with a high concentration of atazanavir and epinephrine and then mixed with purified heterotrimeric Gs protein and Nb35 to form a stable complex. The structure of the β_1_AR-Gs-atazanavir was determined at 3.13 Å **(Extended Data Fig. 6 and Extended Data Table 1)**. The clear electron densities enable the modeling of atazanavir, epinephrine and the majority of the receptor, except for the flexible ICL3 (residues 261-315) **(Fig. 4a)**. The overall structure of the β_1_AR-Gs-atazanavir is similar to previously reported β_1_AR-Gs structures^25,26^, with TM6 adopting an outward-displaced conformation **(Extended Data Fig. 7a)**. A well-defined density for atazanavir is observed in the shallow pocket between TM6 and TM7, with its binding pose slightly shifted towards TM7 compared with GPR119 **(Fig. 4e)**. Similar to the mutagenesis studies in GPR119, increasing potential steric clashes by mutating I328^6.39^ to W in the β_1_AR also impairs the function of atazanavir **(Fig. 4b,c)**, suggesting that this position may be conserved and crucial for atazanavir binding. Unlike GPR119, the residues F^7.48^ and R^7.55^ on the TM7 of the β_1_AR appear to directly interact with atazanavir **(Fig. 4b)**. Consistent with these structural observations, the F372^7.48^A/R379^7.55^D double mutation (β_1_AR-2mut) abolishes the function of atazanavir while leaving the activation by isoprenaline unaffected **(Fig. 4d)**. These structural comparisons underscore the dynamic and adaptable characteristics of atazanavir’s binding, highlighting its ability to engage distinct GPCR systems effectively.

**Figure 4.**
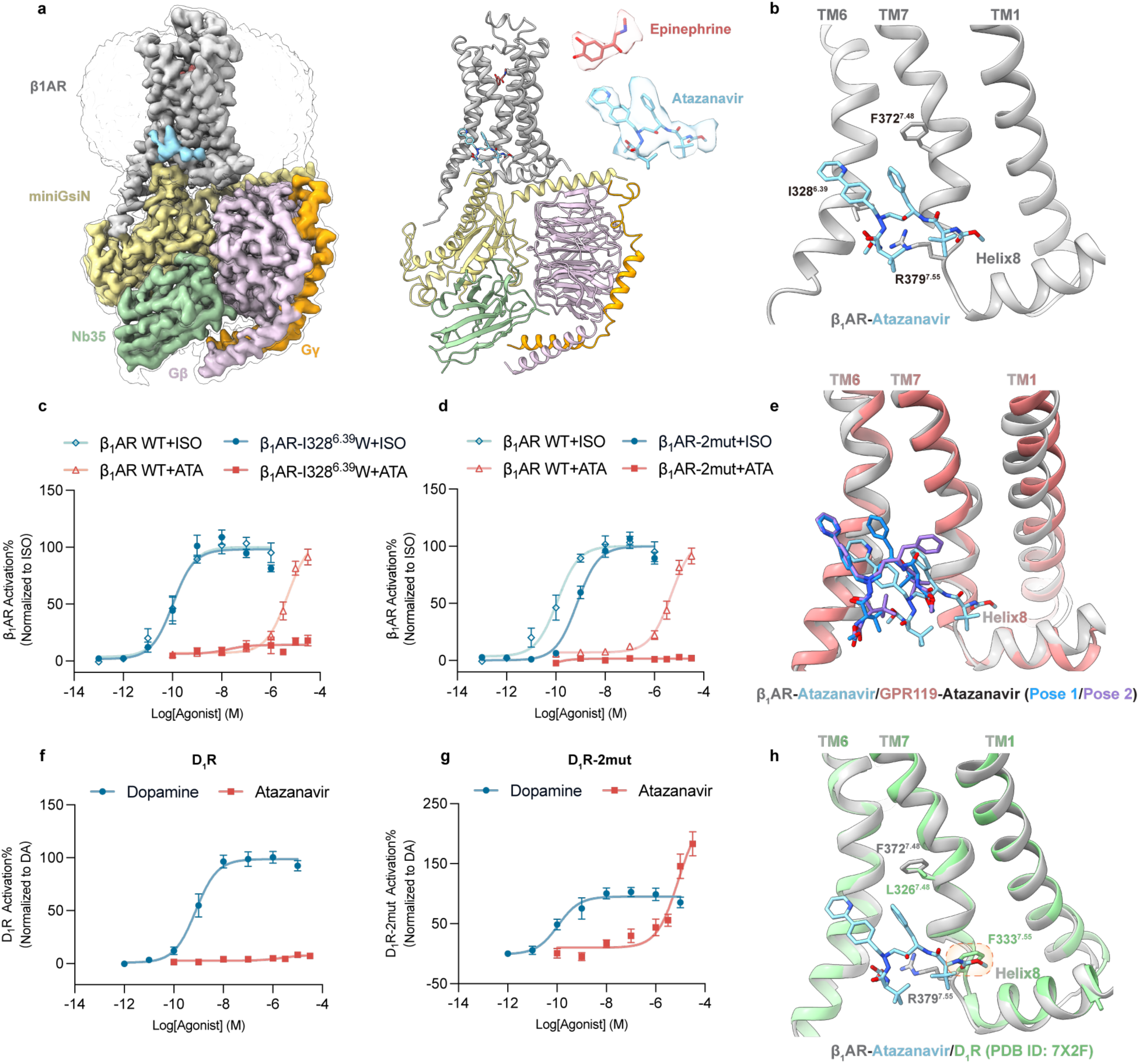
Cryo-EM structure of atazanavir-bound β_1_AR-Gs complex and comparison of atazanavir’s binding poses in different receptors. **a,** Overall structure of atazanavir bound β_1_AR-Gs-Nb35 complex. From left to right: electron density map, coordinates model and the zoom-in view of the electron densities for the ligands. β_1_AR, silver; Gαs, yellow; Gβ, violet; Gγ, orange; Nb35, light green; epinephrine, indian red; atazanavir, sky blue. **b,** Intramolecular interactions between atazanavir and residues in the binding pocket of β_1_AR. **c-d,** β_1_AR-I328^6.39^W (**c**) and β_1_AR-2mut (**d**) mutants can’t be activated by atazanavir in cAMP GloSensor assay. Data are given as Mean ± S.E.M. of 6-7 independent replicates. **e,** The atazanavir binding pose in the β_1_AR slightly shifts towards TM7 compared with GPR119. **f-g,** Atazanavir is incapable of activating wild-type D_1_R (**f**), but can activate D_1_R-2mut with L326^7.48^ F and F333^7.55^ R mutation (**g**) in cAMP GloSensor assay. Data are given as Mean ± S.E.M. of 8-10 independent replicates. **h,** Residues in the atazanavir’s binding pocket are different between D_1_R (pale green) and β_1_AR (silver).

The structural information of atazanavir bound to β_1_AR also guided the design of gain-of-function (GoF) mutation. As previously mentioned, D_1_R does not respond to atazanavir **(Fig. 4f)**. Sequence and structure alignments reveal that the residues involved in atazanavir’s binding differ between D_1_R and β_1_AR, with F333^7.55^ in D_1_R sterically clashing with atazanavir’s carbamate group **(Fig. 4h).** As expected, when two residues in D_1_R (L326^7.48^ F /F333^7.55^ R) were mutated to mimic the atazanavir binding pocket in β_1_AR, the resulting mutant receptor (D_1_R-2mut) could be activated by atazanavir while maintaining its ability to respond to othorsteric agonist dopamine **(Fig. 4g)**.

### Sustained signaling of atazanavir on Family A GPCRs

In a cAMP GloSensor assay, we observed surprising differences in the activation profiles of atazanavir and AR231453 on GPR119. The AR231453-induced cAMP accumulation peaks at approximately 5 minutes and subsequently declines, whereas the atazanavir-induced cAMP accumulation is maintained for over 90 minutes **(Fig. 5a)**. It is noteworthy that AR231453 was previously reported to exhibit wash-resistant sustained activation of GPR119 in the presence of the PDE inhibitor IBMX^27^, likely due to its slow dissociation. However, under our experimental conditions, AR231453 does not demonstrate this sustained activation. G protein-biased signaling may explain the unique signaling profile of atazanavir, as β-arrestin recruitment is generally believed to inhibit G protein signaling^28^. However, as previously mentioned, NanoBiT-based β-arrestin recruitment assay demonstrates that atazanavir can trigger β-arrestin recruitment for GPR119 **(Fig. 1e)**. These results suggest that atazanavir-bound GPR119 can trigger the Gs protein signaling for more than 90 minutes while simultaneously recruiting β-arrestin, indicating a novel mechanism where G protein and β-arrestin signaling coexist without mutual inhibition. Moreover, atazanavir’s ability to induce sustained activation is not limited to GPR119. Similar effects are observed in other GPCRs, including β_1_AR **(Fig. 5a)**, β_2_AR, M_3_R, α_1_AR and D_1_R-2mut **(Extended Data Fig. 8a)**.

**Figure 5.**
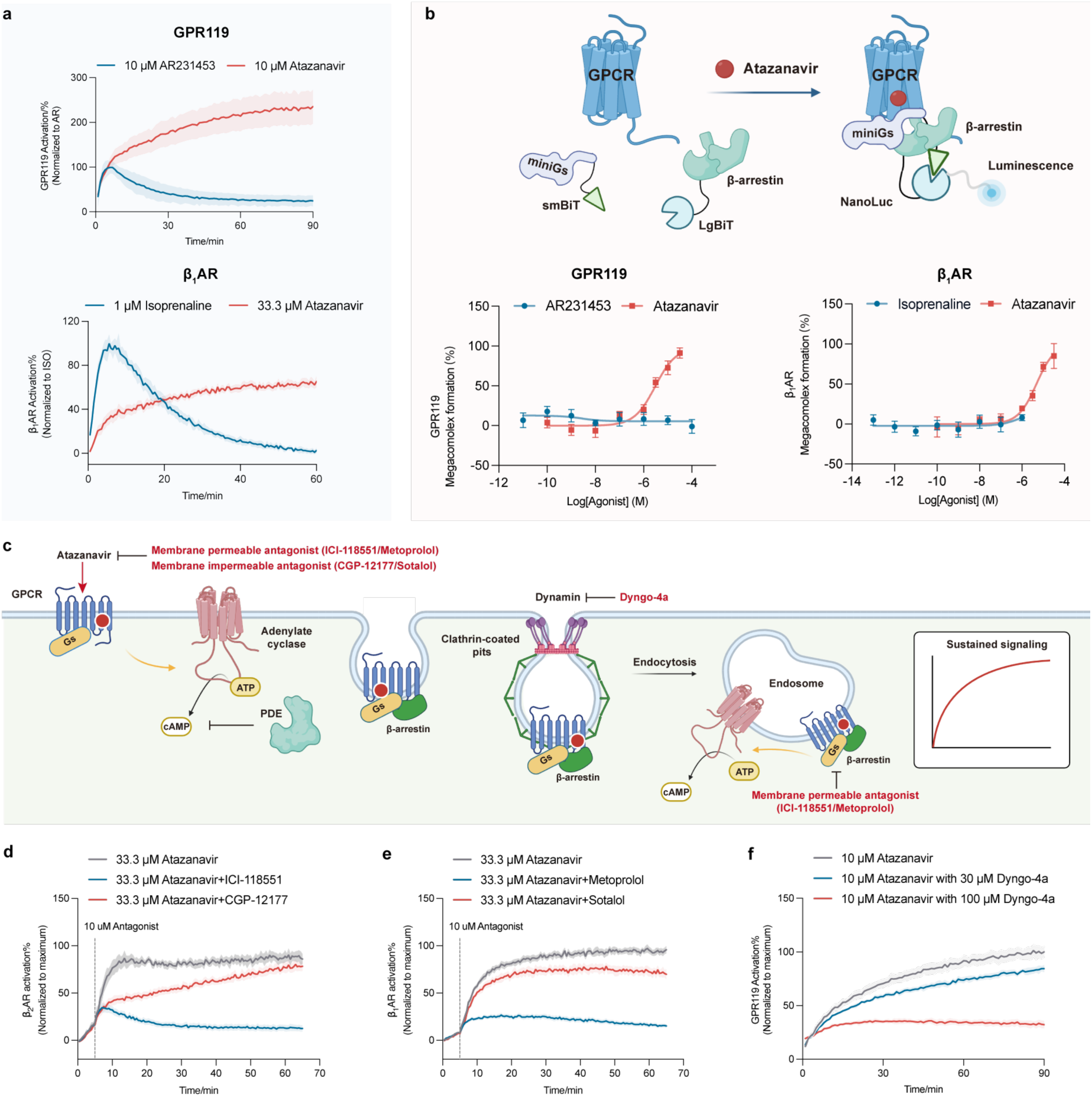
Atazanavir mediates sustained G protein signaling in internalized compartments by forming GPCR - Gs protein - β-arrestin megacomplexes. **a,** Atazanavir induces sustained G protein signaling in GPR119 and β_1_AR in cAMP GloSensor assay, while AR231453 or isoprenaline does not. Data are given as Mean ± S.E.M. of 8 independent replicates. **b,** NanoBiT-based miniGs-β-arrestin co-localization assay suggests the proximity of miniGs and β-arrestin when GPR119 and β_1_AR are activated by atazanavir. Data are given as Mean ± S.E.M. of 6-9 independent samples. **c,** Proposed mechanism of atazanavir-simulated sustained activation of Family A GPCRs. Atazanavir-stimulated receptor mediates sustained endosomal G protein signaling, during which the β-arrestin and G protein are spatially close to each other. The figure was created with BioRender. **d-e,** Atazanavir-induced sustained G protein signaling of β_1_AR (**d**) and β_2_AR (**e**) can only be totally inhibited by membrane-permeable antagonists (ICI-118551/metoprolol) rather than membrane-impermeable antagonists (CGP-12177/sotalol) in cAMP GloSensor assay. Data are given as Mean ± S.E.M. of 6 independent replicates. **f,** Dyngo-4a inhibits atazanavir-stimulated GPR119 activation. Data are given as Mean ± S.E.M. of 3-4 independent replicates.

Sustained G protein signaling has been reported for several Class B GPCRs^6–8^, whose carboxyl-termini exhibit high affinity to β-arrestin and are capable of forming GPCR - G protein - β-arrestin magecomplexes through the tail-engagement of β-arrestin^29,30^. In another case, replacing the carboxyl terminus of β_2_AR (Class A) with that of V_2_R (Class B) results in a β_2_V_2_R chimeric receptor capable of forming a GPCR - G protein - β-arrestin magacomplex and exhibiting sustained G protein signaling^31,32^. To investigate whether atazanavir can mediate sustained signaling by promoting the formation of similar magecomplexes, we employed a NanoBiT-based colocalization assay. SmBiT was fused to miniG and LgBiT was fused to β-arrestin1 to detect the formation of GPCR - Gs protein - β-arrestin megacomplexes **(Fig. 5b)**. The results indicate that miniG and β-arrestin are in close proximity during the activation process, suggesting that atazanavir promotes the formation of megacomplex for GPR119, as well as other GPCRs responsive to atazanavir, including β_1_AR **(Fig. 5b)**, β_2_AR, M_3_R, α_1_AR, μOR and even D_1_R-2mut **(Extended Data Fig. 8b,c)**. In contrast, orthosteric agonists of these receptors result in unchanged or even decreased luminescence signals in the same colocalization assay. This observation is consistent with the canonical signaling paradigm, where G protein and β-arrestin compete for binding to the same intracellular cavity of the receptor.

Previous investigations into the GPCR - G protein - β-arrestin megacomplex have shown that sustained signaling originates from internalized compartments^31^. To explore whether atazanavir-induced sustained signaling similarly arises from internalized compartments, β_2_AR- overexpressing cells were firstly stimulated by atazanavir or isoprenaline for 5 minutes. Subsequently, the cells were treated with either a membrane-impermeable β_2_AR antagonist (CGP- 12177) to inhibit Gs signaling at the cell membrane or with a membrane-permeable β_2_AR antagonist (ICI-118551) to inhibit Gs signaling from both the cell membrane and the internalized compartments **(Fig. 5c).** The results show that for the isoprenaline-induced Gs signaling of β_2_AR, both antagonists suppress the Gs signal to a similar extent **(Extended Data Fig. 8d)**. However, in the case of atazanavir-activated β_2_AR, ICI-118551 fully blocks Gs signaling, while CGP-12177 only partially inhibits it **(Fig. 5d).** A similar phenomenon is observed in β_1_AR. A membrane-permeable β_1_AR antagonist (metoprolol) completely inhibits atazanavir-induced Gs signaling, whereas a membrane-impermeable β_1_AR antagonist (sotalol) exhibits only partial inhibition **(Fig. 5e and Extended Data Fig. 8e)**. These results suggest that a significant portion of cAMP stimulated by atazanavir originates from internalized compartments through the formation of intracellular megacomplexes^31^.

We further investigated whether inhibiting β-arrestin-mediated internalization would suppress the prolonged cAMP accumulation, given that sustained signaling from intracellular compartments depends on the β-arrestin-induced internalization. The presence of Dyngo-4a, an inhibitor for dynamin, significantly impairs atazanavir-stimulated GPR119 activation **(Fig. 5f).** Taken together, these results indicate that atazanavir-induced prolonged GPCR activation is correlated with receptor internalization.

## Discussion

The therapeutic potential of GPCR ligands has led to intense interest in ligand screening targeting GPCRs. The screening process can be conducted either virtually or experimentally. Virtual screening enables rapid screening of millions of compounds in a short period of time^33^, while experimental screening has the potential to identify functional modulators that target previously unexplored binding sites^34^. This work introduces a new approach for the experimental screening of GPCR agonists by converting the activation signals of target GPCRs to yeast growth events. The method leverages a protein-fragment complementation strategy that takes advantage of the coupling of GPCR and G protein upon receptor activation, utilizing split Trp1p as a reporter protein. We present the establishment of the SPS system for GPR119 and β_2_AR in this work. However, the strategy is likely broadly applicable to other GPCRs due to the conserved interaction pattern between GPCRs and G proteins. It is important to mention that false-positive hits may occur for compounds like indole or tryptophan, which enable yeast to bypass the need for Trp1p. But these false positives could easily be distinguished from true signals based on their chemical structures.

As a proof of concept, we applied the SPS method for medium-scale screening and identified atazanavir as a novel positive allosteric modulator for GPR119, a potential target for the treatment of type 2 diabetes. Surprisingly, we discovered that atazanavir is also capable of activating many other Family A GPCRs that couple to different G proteins, including β_1_AR, β_2_AR, M_3_R, α_1_AR and μOR. The versatility of atazanavir makes it a pan-PAM for Family A GPCRs. The structures of the atazanavir-bound GPR119-Gs complex and β_1_AR-Gs complex reveal that atazanavir binds to the membrane-facing cytoplasmic side of TM6/7. The atazanavir binding pose is slightly different between GPR119 and β_1_AR, likely attributable to variations in surrounding residues. We hypothesize that atazanavir interacts with other GPCRs at the same shallow pocket between TM6 and TM7, albeit with slight conformational adjustments. The broad activation of diverse GPCRs by atazanavir may account for its various side effects^35^. Moreover, atazanavir’s pan-activity across diverse GPCRs positions it as a valuable tool for investigating Family A GPCR functions, such as the function of orphan GPCRs.

The most intriguing feature of atazanavir is its sustained activation profile for GPR119 and other Family A GPCRs. This phenomenon is demonstrated to arise from the atazanavir-induced formation of a noncanonical receptor-β-arrestin complex that remains compatible with G protein coupling and occurs within intracellular compartments. In our companion work, we solved the cryo-EM structure of the atazanavir-induced GPCR - G protein - β-arrestin megacomplex, thereby fully revealing the molecular mechanism by which atazanavir stabilizes this complex and mediates sustained G protein signaling. Traditionally, β-arrestin recruitment is thought to terminate G protein signaling, and the typical approaches to prolong G protein signaling involve diminishing the β-arrestin recruitment activity to develop “G protein biased agonists”^36^. Our results, however, represent a new mechanism, offering a novel positive allosteric modulator for Family A GPCRs as a general starting point to achieve sustained G protein signaling. This approach may offer pharmacological advantages in scenarios where prolonged G protein signaling is preferred. However, further modifications are necessary to improve the affinity and selectivity of atazanavir for specific GPCR targets, ensuring more precise therapeutic applications.

## Methods

### pYD-GPCR-SPS plasmid construction

In order to use Trp1p as a reporter protein of Survival Pressure Selection, the TRP1 auxotrophic maker on the pYD1 plasmid was exchanged into a LEU2 auxotrophic marker. The resulting pYD1-Leu plasmid was used for SPS system establishment. The cDNA of human β_2_AR(Δ343-413) and miniGs were subcloned into pYD-Leu plasmid at the upstream and downstream of *GAL1,10* bidirectional promoter, respectively, for simultaneous expression under the induction of galactose in the medium. The N-terminus sequence of Trp1p (Trp1p^NTD^) was amplified and flanked by the homogenous sequence of miniGs for DNA insertion, while a GGS flexible linker was added between miniGs and both terminus of Trp1p^NTD^. The C-terminus sequence of trp1p (Trp1p^CTD^) was amplified and flanked by the homogenous sequence of β_2_AR together with a GGSGG linker for C-terminus fusion. Constructed plasmids (name pYD-SPS- β_2_AR) with different Trp1p split sites were electroporated into EBY100 for agonist-dependent yeast growth validation. The pYD-SPS-GPR119 plasmids were constructed in the same way.

### Yeast electroporation

A single colony of EBY100 was inoculated into the YPD medium and cultured at 30 °C overnight for recovery. On the next day, the saturated yeast culture was diluted to OD_600_ = 0.2 and continuously cultivated to OD 600 = 1.5. For competent EBY100 yeast cell preparation, yeast cells were spun down at 3500 × g for 5 min to remove the medium. The pellet was washed with pre-chilled MQ water and then resuspended into 1 M Sorbitol with 10 mM Tris-HCl pH=8, and 10 mM LiAc and incubated at 30 °C for 30 min. 10 mM DTT was directly added into the yeast cells for another 15 min incubation to get the competent EBY100 cells. After incubation, competent EBY100 cells were collected and washed with pre-chilled MQ water again. Finally, the cell pellet was resuspended in pre-chilled MQ water and was ready for electroporation. For electroporation, usually, 1 μg plasmid DNA was mixed with 50 μl competent EBY100 cells in an ice-cold cuvette. The electroporation is performed with the following parameters: 500 Voltage square pulse, 15 ms with one pulse only. Electroporated yeast was transferred into a sterile tube with pre-warmed YPD medium for recovery at 30 °C for 1 min. After recovery, yeast was spun down and plated onto - LEU plates to separate single colonies.

### HTS screening of GPR119 agonists

The selected drug libraries include the Selleck FDA-approved Drug Library, Selleck Preclinical & Clinical Compound Cherry Pick Library, TargetMol Natural Compound Library, MCE Natural Product Library, MetaSci Human Metabolite Library, MCE GPCR/G protein Compound Library. The assays were performed in 384 well plates. EBY100-SPS-GPR119 single colony was inoculated into -LEU/glucose medium for yeast growth and diluted into -TRP/- LEU/galactose medium for screening. Compound libraries are provided in 384-well plates and the compounds were transferred to sterile transparent 384-well culture plates by Echo500 at a volume of 50 nl per well. Yeast was then inoculated into these culture plates at a volume of 40 μl per well. OD600 of each well was measured by a multi-plate reader and used as the baseline. Reported agonists of GPR119 were also added into several wells as positive control. Yeast plates were incubated under 25 °C and 85% humidity and OD 600 of each well was measured daily with the same sequence as the first day. Data were processed in MATLAB to determine OD 600 change of each well by subtracting the baseline value from it.

### Expression and purification of Gs heterotrimer, Gβ1γ2 heterodimer, scFv16 and nanobody-35

The expression and purification of Gs heterotrimer and Gβ1γ2 heterodimer were performed following previously described protocols^22^. In brief, Gs heterotrimer and Gβ1γ2 heterodimer were expressed in *Trichoplusia ni* (Hi5) insect cells by co-infecting Hi5 cells with baculoviruses encoding the bovine Gαs subunit and rat Gβ1-bovine Gγ2 subunits or with baculoviruses encoding Gβ1γ2 subunits only. The Gβ1γ2 subunits were tagged with an N-terminal 6×His tag followed by a 3C protease cleavable site (LEVLFQGP). After expression, the target proteins were purified using nickel affinity chromatography. HRV-3C protease, prepared in-house, was added to the eluted protein, and the mixture was dialyzed to remove excess imidazole from the elution buffer. To remove HRV-3C protease and any contaminating proteins, the sample was subjected to a second Ni-NTA affinity chromatography, during which the flow-through was collected.

scFv16 was expressed and purified as described previously^37^. Briefly, scFv16 was expressed in secreted form from Hi5 insect cells and subsequently purified using nickel affinity chromatography followed by size exclusion chromatography on a Superdex 75 increase 10/300 column (Cytiva).

Nanobody-35 (Nb35) was expressed in the *E. coli* strain BL21(DE3) and purified according to previously described methods^22^. The purification process involved nickel affinity chromatography, followed by size exclusion chromatography on a Superdex 75 increase 10/300 column (Cytiva).

### GPR119-miniGsiN Construct Design and Expression

Human GPR119 with an N-terminal HA signal sequence followed by a FLAG tag was inserted into the pcDNA 3.1 vector. The miniGsiN protein^38^ was fused to the GPR119 C-terminus connected by a 3C protease recognition site (LEVLFQGP), His tag, and flexible glycine/serine linkers. The constructed plasmid was transfected into Expi293F cells with PEI, and the cells were maintained in SMM 293-TI (Sino biological Inc) medium at 37°C. Cells were collected 48 hours after transfection and stored at −80°C for further use.

### Purification of GPR119 Complex

GPR119-miniGsiN was solubilized in solubilization buffer (20 mM HEPES, pH 7.5, 100 mM NaCl, 2 mM MgCl_2_, 1% DDM, and 0.02% CHS). Excess agonist and a 1.2 × molar ratio of purified Gβ1γ2 heterodimer, Nb35, scFv16, were incubated with the receptor for 2 hours during solubilization. The complex was purified by M1 Anti-FLAG affinity chromatography (Sigma) with detergent exchange to 0.01% L-MNG (Anatrace), 0.002% CHS, followed by size exclusion chromatography on a Superdex 200 increase 10/300 column in a final buffer comprised of 20 mM HEPES, pH 7.5, 100 mM NaCl, 0.002% LMNG, 0.0004% CHS, and 50 μM agonist. The purified complex was then concentrated to 5 mg/ml using a 100 kDa MWCO concentrator, flash-frozen in liquid nitrogen, and stored at −80°C.

### Expression and Purification of β_1_AR Complex

The human β_1_AR was expressed in *Spodoptera frugiperda* (SF9) insect cells with recombinant baculovirus (Bac-to-Bac expression systems) for 48 h at 27 °C. The β_1_AR was solubilized from the cell membrane in solubilization buffer (20 mM HEPES, pH 7.5, 100 mM NaCl, 1% LMNG and 0.02% CHS) and purified by M1 Anti-FLAG affinity chromatography (Sigma) followed with a size exclusion chromatography on a Superdex 200 increase 10/300 column in a final buffer comprised of 20 mM HEPES, pH 7.5, 100 mM NaCl, 0.01% LMNG, 0.002% CHS. The purified β_1_AR was mixed with agonists and incubated with a 1.2 × molar excess of purified Gs heterotrimer and Nb35 for 2 hours at room temperature. A final size exclusion chromatography purification was performed to remove excess Gs heterotrimer and Nb35. The size exclusion chromatography buffer was comprised of 20 mM HEPES, pH 7.5, 100 mM NaCl, 0.002% LMNG, 0.0004% CHS and 50 μM agonist. The purified complex was then concentrated to 8 mg/ml using a 100 kDa MWCO concentrator, flash frozen in liquid nitrogen and stored at −80°C.

### Cryo-EM sample preparation and data collection

For the cryo sample preparation of GPR119-miniGsiN complexes and β_1_AR-Gs complexes, 4 µl of protein was loaded onto glow-discharged holey carbon grids (Quantifoil Au R1.2/1.3, 200 mesh or 400 mesh). The grids were blotted for 4.5 seconds and flash-frozen in liquid ethane using a Vitrobot (Mark IV, Thermo Fisher Scientific). Cryo-EM data were collected using 300 kV Titan Krios equipped with Gatan K3 Summit detector or Falcon 4 electron detector at Shuimu Biosciences. Movie stacks were collected with a defocus range from −1.3 µm to −1.6 µm with a total dose of about 50 e-/Å^2^ using EPU data acquisition software in super resolution mode. Each stack was exposed for 2.56 seconds, and 32 frames per micrograph were aligned by MotionCor2^39^ and binned to a pixel size of 0.86 Å (GPR119-Gs-AR231453 and GPR119-Gs-atazanavir) or 1.08 Å (β_1_AR-Gs-atazanavir).

### Cryo-EM data processing and structure determination

Before being imported into cryoSPARC^40^, dose-weighted micrographs were manually picked using the kdisplay (https://github.com/phoeningo/ksoft.git).

For the GPR119-AR231453 complex, 2,817,323 particles were picked from 5255 micrographs by blob-picker. Two rounds of 2D classification were performed, and particles exhibiting features of a GPCR-G protein complex were selected to generate initial models through ab-initio reconstruction. Subsequently, multiple rounds of heterogeneous refinement were conducted, and 485,493 particles were re-extracted for further calculation. After heterogeneous refinement, non-uniform refinement and local refinement, the final cryo-EM map was obtained with a resolution of 2.98 Å **(Extended Data Fig. 2)**.

For the GPR119-Gs-atazanavir complex, 3,109,923 particles were picked from 4482 micrographs using the templates from the GPR119-Gs-AR231453 dataset. Obvious junk particles were removed after one round of 2D classification. The remaining particles were further classified through heterogeneous refinement using the cryo-EM map of GPR119-Gs-AR231453 as a reference. After heterogeneous refinement, 1,268,255 particles were selected and re-extracted for additional 2D classification. From these, 257,037 particles with clear features of a GPCR-G protein complex were chosen for ab initial reconstruction. All 1,268,255 particles were then subjected to non-uniform refinement by using the initial map as a reference, followed by iterative 3D classifications to identify the density of atazanavir. Ultimately, 106,271 particles were selected and refined to achieve a global resolution of 3.33 Å, with the density of atazanavir clearly defined, enabling unambitious model building **(Extended Data Fig. 3)**.

For the β_1_AR-Gs-atazanavir complex, a total of 2,759,995 particles were initially auto-picked from 1,591 micrographs. After iterative 2D classification, particles with clear features of a GPCR-G protein complex were selected as templates for subsequent particle picking. Following the removal of poor-quality micrographs, 978,700 particles were picked and extracted based on these templates. These 978,700 particles were subjected to one round of heterogeneous refinement using the canonical cryo-EM map of a GPCR-G protein complex as a reference. After classification, 289,920 particles were selected for ab-initio reconstruction to generate the initial map of the β_1_AR-Gs-atazanavir complex. Additionally, 1,276,345 particles were picked and extracted from another 1,248 micrographs. Combining these particles with the previously selected 978,700 particles, a total of 2,255,045 particles were subjected to heterogeneous refinement using the initial map as a reference. A total of 743,660 particles were selected and re-extracted for heterogeneous refinement and non-uniform refinement. To improve the occupancy of atazanavir, receptor-focused 3D classifications were performed. Finally, 132,798 particles were selected to generate a new initial map through *Ab initial*. The clear density of atazanavir around TM6 and TM7 of the receptor can be observed from the initial map. Then, the particles were refined against the initial map and achieved a final resolution of 3.13 Å. The density of atazanavir is clear enough for model building **(Extended Data Fig. 6)**.

Coordinates and restraints of atazanavir and AR231453 were generated using eLBOW in Phenix1.20.1. For the structures of the GPR119-Gs complex with different ligands, the model of GPR119-Gs-LPC (PDB ID: 7XZ5) was initially docked into the cryo-EM map using UCSF ChimeraX-1.5^41^. This was followed by iterative manual adjustments in COOT0.9.8.3^42^ and real-space refinement in Phenix1.19.2^43^. For the structure of β_1_AR-Gs-atazanavir complex, models of human β_1_AR (PDB ID:7BTS) and Gs-Nb35 (PDB ID: 3SN6) were first docked into the cryo-EM map using UCSF ChimeraX-1.5. The subsequent refinement process involved iterative manual adjustments in COOT0.9.8.3^42^ and real-space refinement in Phenix1.19.2^43^. The refinement statistics are provided in Extended Data Table 1. Structural figures were generated using UCSF ChimeraX-1.5.

### cAMP GloSensor assay

Receptors were subcloned into the pcDNA3.1 vector with an HA signal peptide and a FLAG tag at its N-terminus. The day before transfection, CHO-K1 cells were plated into 6-well plates in DMEM/F12 medium containing 10% FBS at a density of 2 ×10^5^ cells per ml. After overnight growth, cells were co-transfected with plasmid mixtures of receptor constructs and cAMP biosensor GloSensor-22F (Promega). For Gq coupled receptors, a chimeric Gsq protein, which was constructed by replacing the C-terminus of Gs to that of Gq^44^, was also transfected into the cells. After 24 hours, cells were digested by 0.05% trypsin and resuspended with HBSS containing 20 mM HEPES pH=7.5 and 2% GloSensor cAMP Reagent (Promega) and plated into 96-well plates at a volume of 100 μl per well. Cells were incubated at 37 °C and room temperature for 2 hours before measurement. The basal luminescence of each well was determined, and varying concentrations of agonist were added into each well to detect cAMP accumulation. The luminescence signal of each well is measured every half minute. Data were processed by GraphPad Prism 9.0 (GraphPad LLC, CA) and shown as Mean ± SEM.

### NanoBiT-based G protein and β-arrestin recruitment assay

The SmBiT fragment was fused to the C-terminus of receptors to generate receptor-SmBiT constructs. The day before transfection, CHO-K1 cells were plated into 6-well plates in DMEM/F12 medium containing 10% FBS at the density of 2 ×10^5^ cells per ml. After overnight growth, cells were co-transfected with plasmid mixtures of receptor-SmBiT and LgBiT-miniG protein or LgBiT-β-arrestin1. After 24 hours, cells were digested by 0.05% trypsin and resuspended with HBSS containing 20 mM HEPES pH=7.5 and 10 μM coelenterazine and plated into 96-well plates at a volume of 100 μl per well. Cells were incubated at room temperature for 1 hour before measurement. The basal luminescence of each well was determined, and varying concentrations of agonists were added into each well to detect luminescence increase. The luminescence signal of each well was measured every half minute, and the data was processed by GraphPad Prism 9.0 (GraphPad LLC, CA) and shown as Mean ± SEM.

### NanoBiT-based miniGs-β-arrestin co-localization assay

The SmBiT fragment was fused to the N-terminus of miniGs to get the construct of SmBiT- miniGs. The day before transfection, CHO-K1 cells were plated into 6-well plates in DMEM/F12 medium containing 10% FBS at the density of 2 ×10^5^ cells per ml. After overnight growth, cells were co-transfected with plasmid mixtures of receptor, SmBiT-miniG protein and LgBiT-β- arrestin1 to investigate the co-localization of miniG protein and β-arrestin1. After 24 hours, cells were digested with 0.05% trypsin resuspended with HBSS containing 20 mM HEPES pH=7.5 and 10 μM coelenterazine and plated into 96-well plates at a volume of 100 μl per well. Cells were incubated at room temperature for 1 hour before measurement. The basal luminescence of each well was determined, and varying concentrations of agonists were added into each well to detect luminescence increase. The luminescence signal of each well was measured every half minute, and the data was processed by GraphPad Prism 9.0 (GraphPad LLC, CA) and shown as Mean ± SEM.

### Whole-cell ELISA assay

Whole-cell expression levels of the wild-type and mutated receptors were determined by whole-cell ELISA assay^45^. Plasmids of different constructs were transfected into CHO-K1 cells the day before measurement and seeded into a 24-well plate. Cells were washed with ice-cold Tris-Buffered Saline (TBS) twice, fixed by 400 μl 4% PFA for 20 min at room temperature, penetrated by 400 μl 0.25% Triton X-100 for 15 min at room temperature and blocked by 500 μl 1% BSA for 1h. Next, cells were incubated with 200 μl HRP-conjuncted anti-DYKDDDDK antibody for 1h at room temperature in the dark. After the removal of antibody, cells were incubated with 200 μl TMB-ELISA substrate solution and then the reaction was stopped by 200 μl stop solution. Absorbance at 450 nm of each well was quantified by a multi-plate reader to reveal the relative expression level of each mutant.

### Flow Cytometry for receptors expression quantification

CHO-K1 cells were seeded in 12-well dishes at a density of 2 × 10^5^ cells per ml medium supplemented with 10% FBS, penicillin and streptomycin 1 day before transfection. For each well, receptors with different mutations or pcDNA3.1 (vehicle) were transfected to determine the expression level. After 1 day, cells are treated with 0.25% Trypsin and resuspended with HBSS containing 20 mM HEPES pH=7.5 to a volume is 300 ul. 10 nM AlexaFlour-488 labelled M1-anti FLAG antibody are mixed with cells for 15 min in dark. After incubation, cells are washed twice with HBSS containing 20 mM HEPES pH=7.5. The fluorescence of each sample is determined by Flow Cytometer at 488 wavelengths. Data was processed in FlowJo (BD Biosciences).

## Data availability

The cryo-EM density maps of GPR119-Gs-AR231453, GPR119-Gs-atazanavir and β_1_AR- Gs-epinephrine-atazanavir have been deposited into the Electron Microscopy Data Bank (EMDB) under accession code EMDB-62871, EMDB-62880 and EMDB-62884, the model coordinates of GPR119-Gs-AR231453, GPR119-Gs-atazanavir and β_1_AR-Gs-epinephrine-atazanavir have been deposited into the Protein Data Bank (PDB) under accession code 9L79, 9L80 and 9L84.

## Acknowledgements

We thank Prof. Brian K. Kobilka for the discussion and suggestions, and Prof. Asuka Inoue for providing the plasmids for the NanoBiT assay. We also thank support from the High Throughput Screening (HTS) Core Facility, Center of Pharmaceutical Technology, Tsinghua University for support with high throughput screening and the Tsinghua University Branch of China National Center for Protein Sciences (Beijing) as well as Shuimu Biosciences for support with cryo-EM data collection. This work was supported by the Tsinghua-Peking Center for Life Sciences, Beijing Frontier Research Center for Biological Structure, Tsinghua University (G.H., X.X. and X.L.), by National Natural Science Foundation of China (Grant 32122041 to X.L. and Grant 32100968 to X.X.), China Postdoctoral Science Foundation 2021M691809 (X.X.) and by Tsinghua University Initiative Scientific Research Program (X.L.).

## Author contributions

G.H. designed, cloned and screened constructs of β_2_AR-SPS and set up the SPS-based drug screening method with help from Q.S.. X.X. initiated the GPR119 project, designed and validated constructs of GPR119-SPS, performed compound screening, and identified atazanavir as a GPR119 agonist with help from G.H.. Q.S. discovered the pan agonist activity of atazanvir together with G.H.. X.X. purified the GPR119-Gs-atazanavir complex. G.H. and Q.S. purified the β_1_AR-Gs-atazanavir complex with help from X.S.. X.X., G.H., Q.S. and S.Z. prepared cryo-EM samples and collected cryo-EM data. G.H., X.X. and X.L. processed the cryo-EM data with help and suggestions from S.Z., F.K., and C.Y.. Q.S., G.H. and X.X. performed the cAMP GloSensor assay, NanoBiT assay and whole cell ELISA assay. Q.S., G.H., X.X. and X.L. wrote the manuscript. X.L. conceived the project and supervised the overall research. All authors contributed to the editing of the manuscript.

## Materials & Correspondence

Correspondence and material requests should be addressed to Xiangyu Liu.

---

**Extended Data Figure 1.**
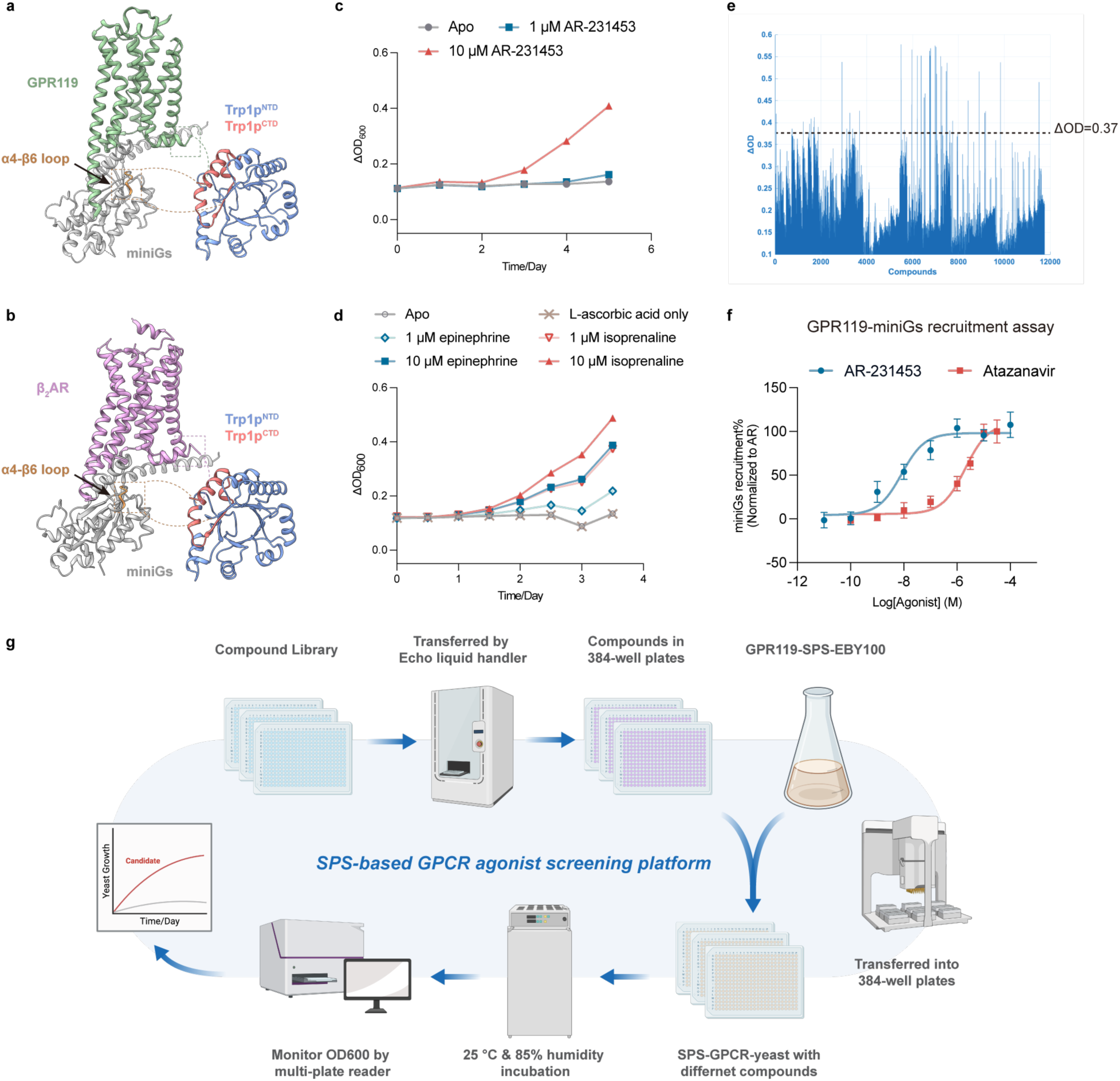
Development of GPCR-SPS system. **a,** The model of GPR119-SPS system. GPR119, green; miniGs, gray; Trp1pNTD, blue; Trp1pCTD, pink. **b,** The model of β_2_AR-SPS system. β_2_AR, violet; miniGs, gray; Trp1pNTD, blue; Trp1pCTD, pink. **c,** The 5-day growth curves of GPR119-SPS yeasts in the presence of 1 μM AR231453, 10 μM AR231453 or without agonist. Data are given as Mean ± S.E.M. of 6 independent replicates. **d,** The 4-day growth curves of β_2_AR-SPS yeasts in the presence of different concentrations of isoprenaline, epinephrine or without ligands. L-ascorbic acid is supplemented to prevent ligand oxidation. Data are given as Mean ± S.E.M. of 6 independent replicates. **e,** Change of optical density (△OD600) of the GPR119-SPS yeasts in around 12,000 different conditions after 6 days of incubation. **f,** Atazanavir activates GPR119 in a NanoBiT-based G protein recruitment assay. Data are given as Mean ± S.E.M. of 8 independent replicates. **g,** Schematic of SPS-based drug screen platform. The figure was created with BioRender.com.

**Extended Data Figure 2.**
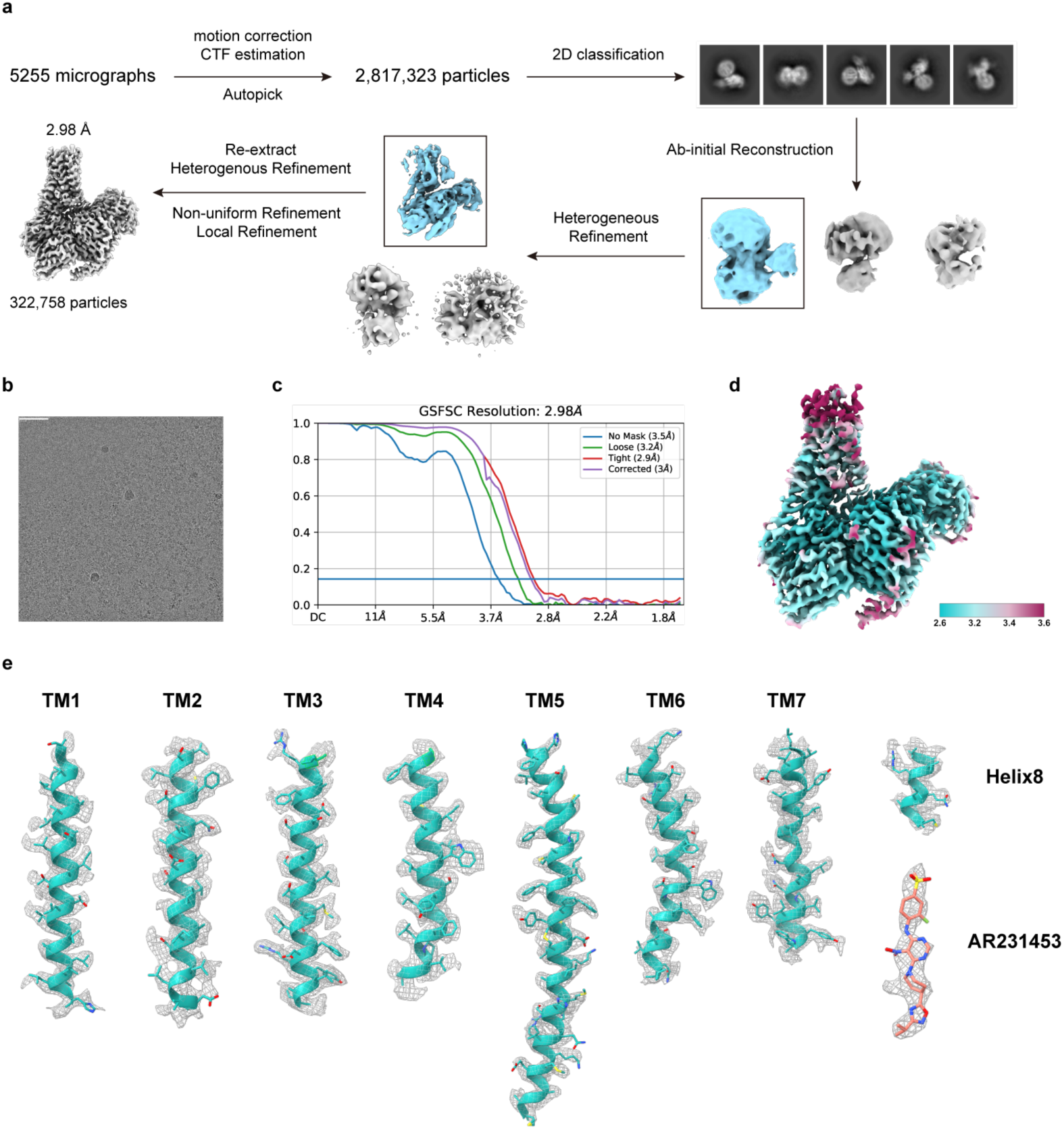
Cryo-EM data processing of GPR119-Gs-AR231453 complex. **a,** Flowchart of cryo-EM data processing of GPR119-Gs-AR231453 complex. **b**, Representative cryo-EM micrograph (scale bar, 50 nm). **c**, Golden Standard Fourier Shell Correlation (GSFCS) curve for the final refinement. **d**, Local resolution of final cryo-EM map. **e**, Cryo-EM map and model of the GPR119-Gs-AR231453 complex. Cryo-EM density map and model are shown for all seven transmembrane α-helices from GPR119 and AR-231453.

**Extended Data Figure 3.**
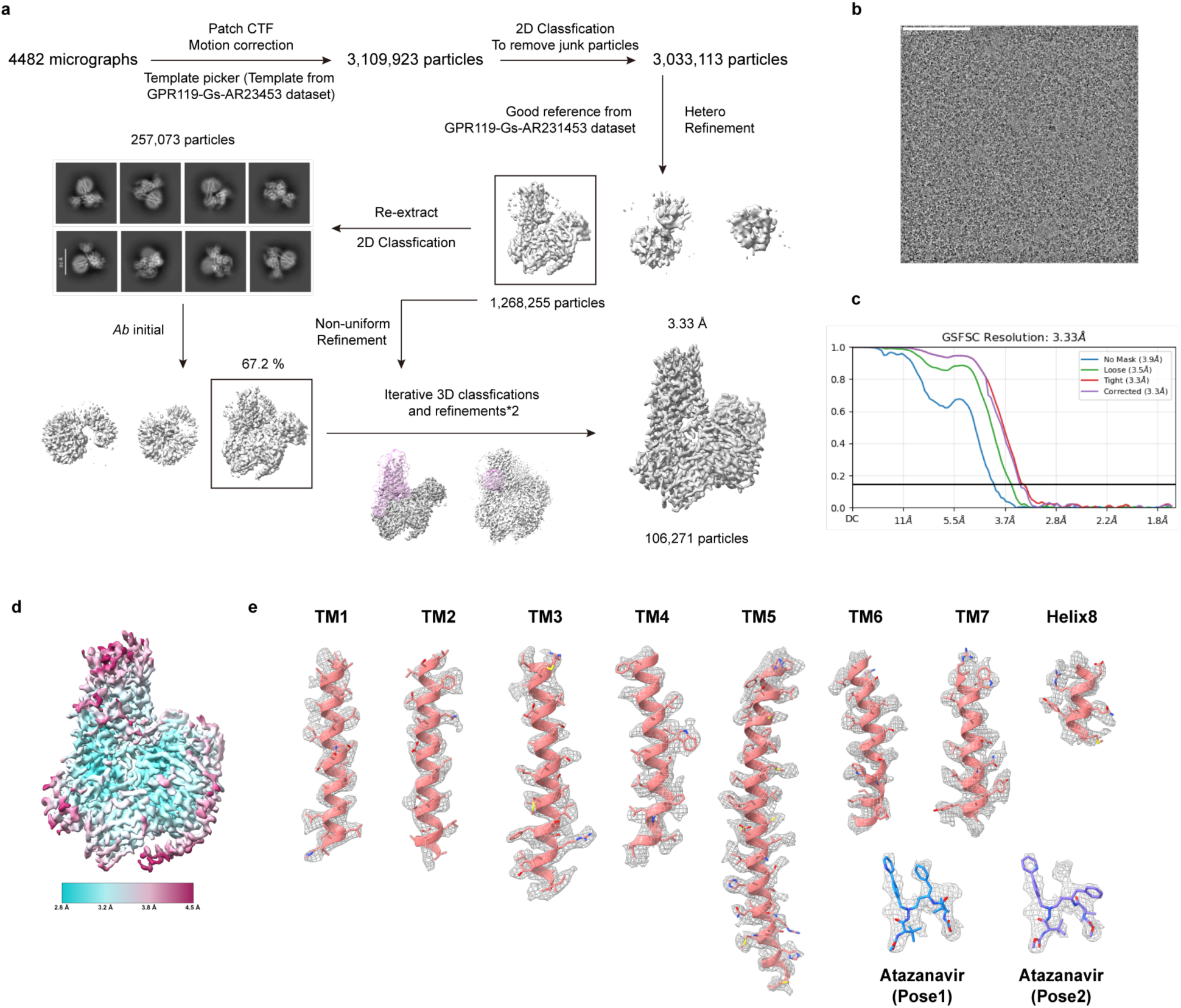
Cryo-EM data processing of GPR119-Gs-atazanavir complex. **a,** Flowchart of cryo-EM data processing of GPR119-Gs-atazanavir complex. **b**, Representative cryo-EM micrograph (scale bar, 50 nm). **c**, Golden Standard Fourier Shell Correlation (GSFCS) curve for the final refinement. **d**, Local resolution of final cryo-EM map. **e**, Cryo-EM map and model of the GPR119-Gs-atazanavir complex. Cryo-EM density map and model are shown for all seven transmembrane α-helices from GPR119 and atazanavir.

**Extended Data Figure 4.**
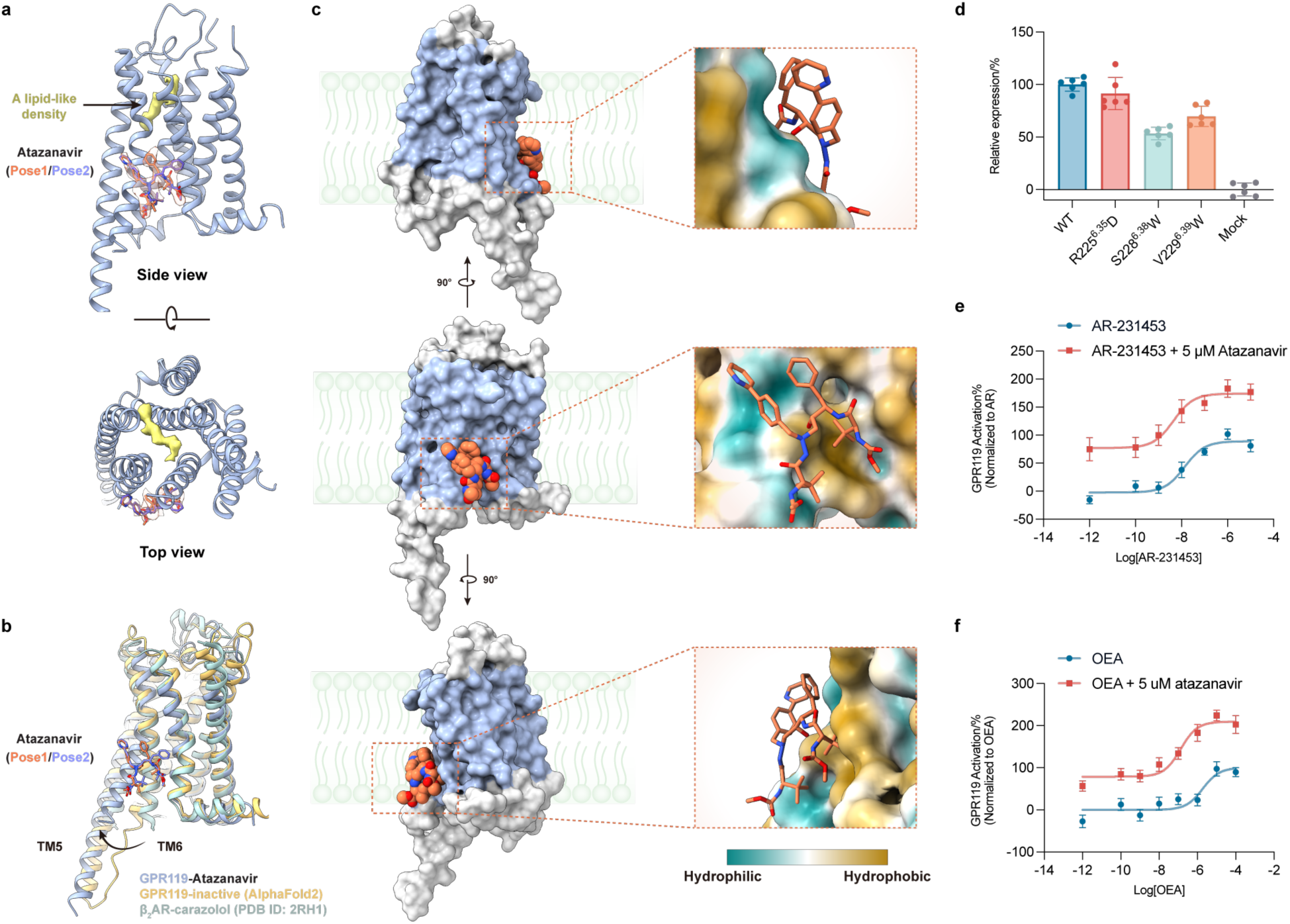
Atazanavir binds to an allosteric site at the transmembrane region of GPR119. **a,** The density in the orthosteric pocket (yellow) shows a similar shape to the endogenous agonist revealed in a recently reported structure of GPR119^30^. **b,** Atazanavir binds to the membrane-facing surface of TM6, which adopts conformational change upon receptor activation. Compared to the structure model of inactive state GPR119 (yellow, predicted by AlphaFold2) and the inactive structure of β_2_AR (light green, PDB ID:2RH1), the intracellular end of TM6 exhibits outward displacement in the active conformation. **c,** Atazanavir is sandwiched between TM6 and the membrane bilayer. The orientation of GPR119 in the lipid bilayer is predicted by the Orientations of Proteins in Membranes (OPM server, https://opm.phar.umich.edu). The predicted transmembrane part of GPR119 is colored in blue. The surface hydrophobicity was calculated using the “molecular lipophilicity potential” (MLP) function in Chimera X. **d,** The expression levels of wild-type and mutated GPR119 were determined by whole cell ELISA. All mutants show comparable expression levels as the wild-type receptor. Data are given as Mean ± S.E.M. of 6 independent replicates. **e-f,** Atazanavir allosterically modulates the agonist activity of AR231453 (**e**) and endogenous agonist OEA (**f**). Data are given as Mean ± S.E.M. of 10-12 independent replicates for e and f.

**Extended Data Figure 5.**
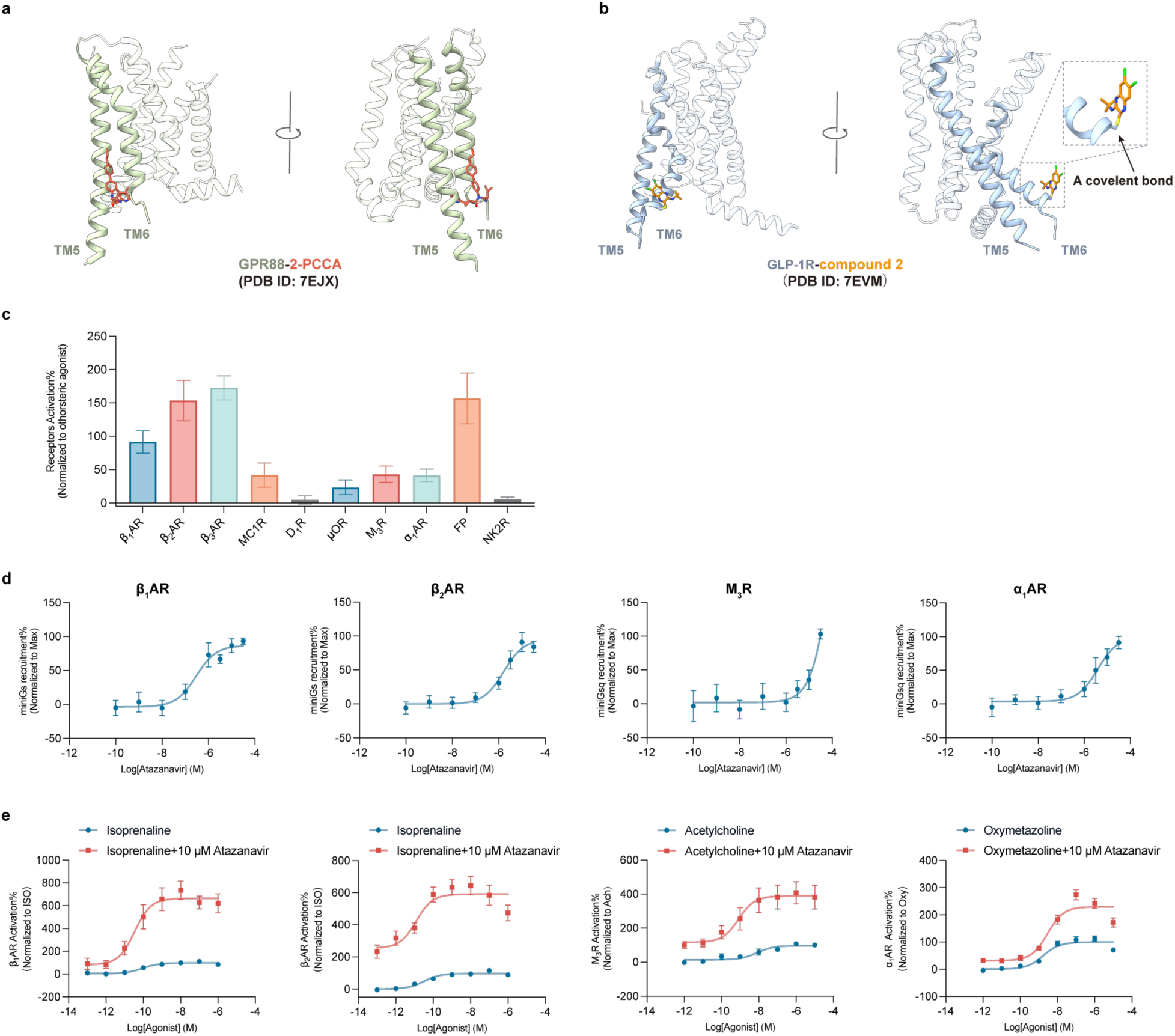
The intracellular end of TM6 represents a general allosteric modulation site for GPCRs. **a-b,** Structures of other allosteric compounds bind to a similar position as atazanavir in GPR119. The structures include 2-PCCA binding to the cleft between TM5 and TM6 of GPR88 (PDB ID: 7EJX, **a**) and compound 2 covalently binding to the surface of TM6 of GLP-1R (PDB ID: 7EVM, **b**). **c,** Atazanavir activates 8 of 10 GPCRs tested in cAMP GloSensor assay or NanoBiT-based G protein recruitment assay. **d,** Atazanavir activates β_1_AR, β_2_AR, M_3_R and α_1_AR in a NanoBiT-based G protein recruitment assay. **e,** Atazanavir allosterically modulates β_1_AR, β_2_AR, M_3_R and α_1_AR, and enhances the EC50 and Emax values of their orthosteric agonists. Data are given as Mean ± S.E.M. of 6-9 independent replicates for c and d.

**Extended Data Figure 6.**
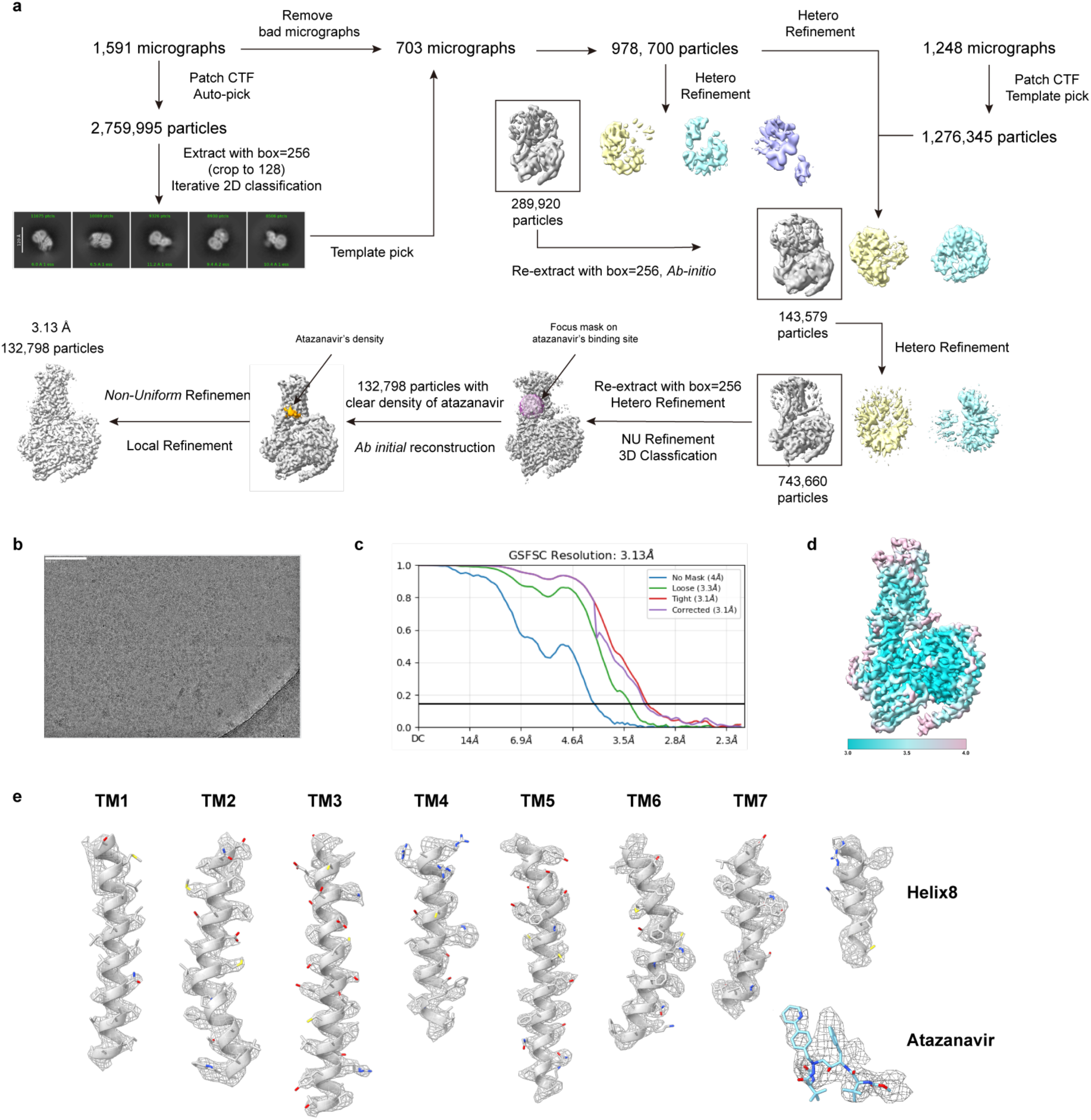
Cryo-EM data processing of β_1_AR-Gs-atazanavir complex. **a,** Flowchart of cryo-EM data processing of β_1_AR-Gs-atazanavir complex. **b,** Representative cryo-EM micrograph (scale bar, 100 nm). **c,** Golden Standard Fourier Shell Correlation (GSFCS) curve for the final refinement. **d,** Local resolution of final cryo-EM map. **e,** Cryo-EM map and model of the β_1_AR-Gs-atazanavir complex. Cryo-EM density map and model are shown for all seven transmembrane α-helices from β_1_AR and atazanavir.

**Extended Data Figure 7.**
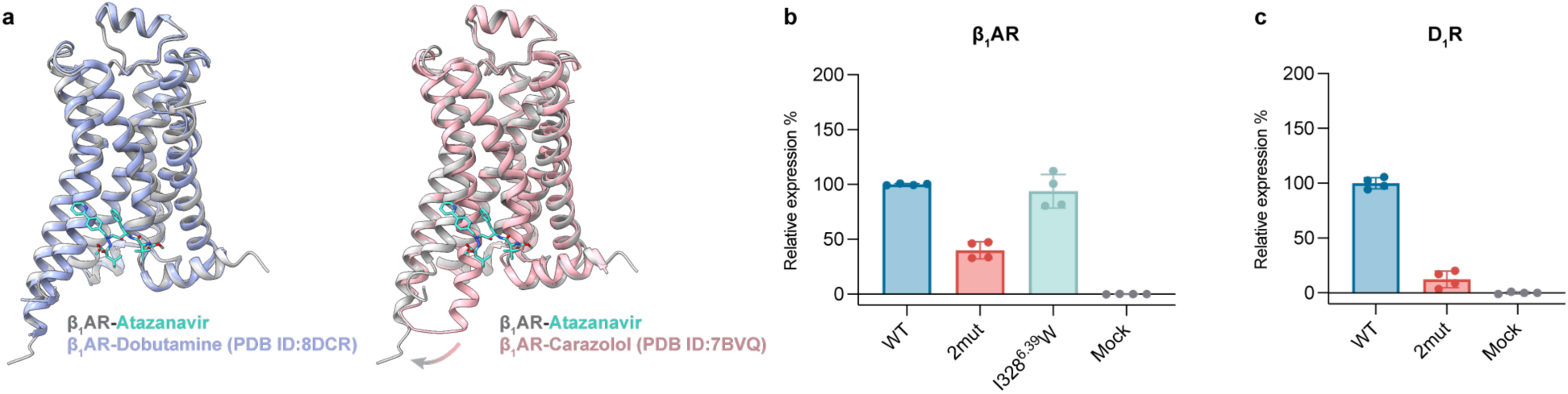
Atazanavir-bound β_1_AR adopts an active conformation. **a,** Comparison of atazanavir-bound active β_1_AR structure (silver) with dobutamine-bound active β_1_AR structure (orchid, PDB ID: 8DCR) and carazolol-bound inactive β_1_AR structure (pink, PDB ID: 7BVQ). **b-c,** The expression levels of wild-type and mutated β_1_AR (**b**) and D_1_R (**c**) were determined by flow cytometry. Data are given as Mean ± S.E.M. of 4 independent replicates.

**Extended Data Figure 8.**
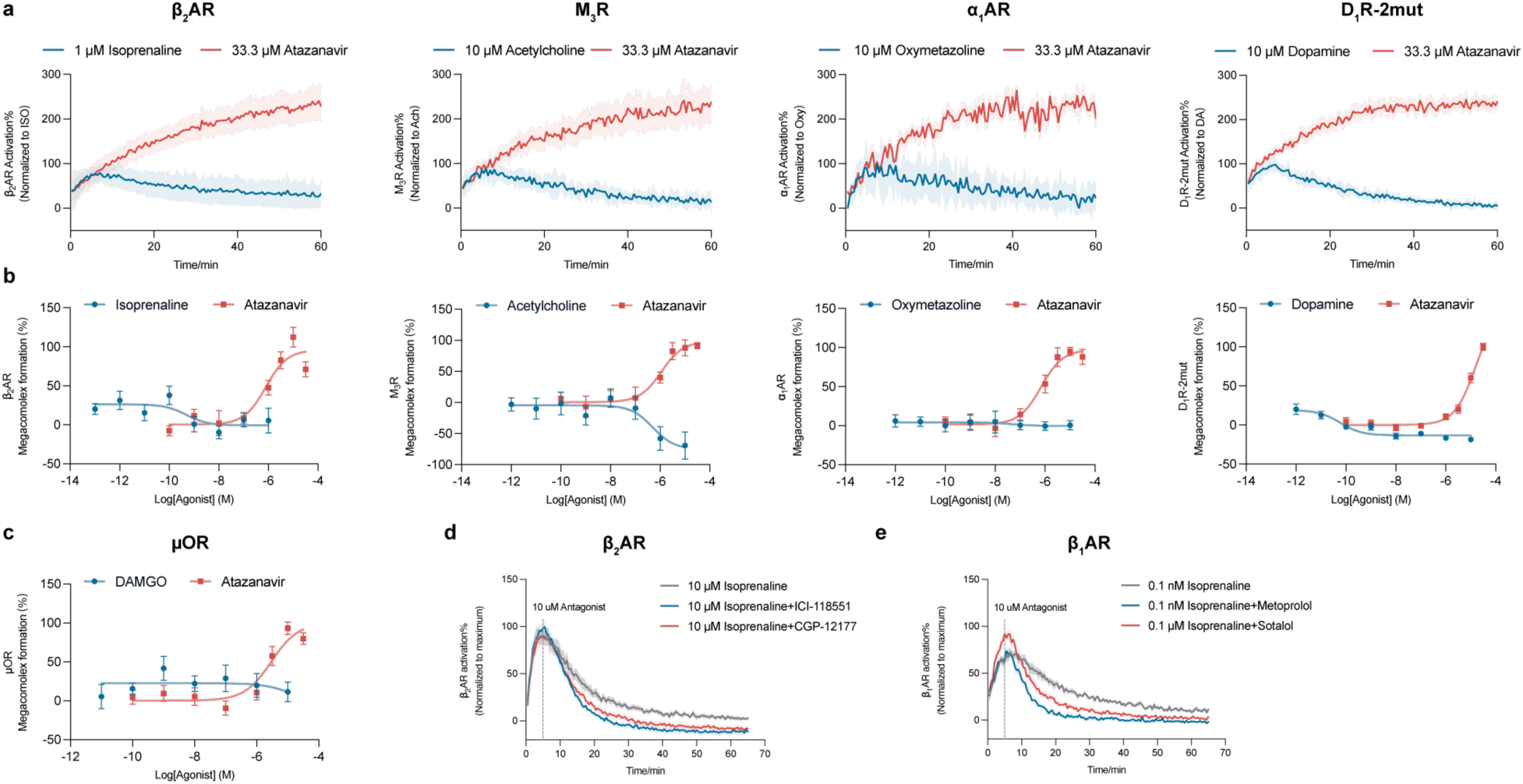
Sustained activation of Family A GPCRs by atazanavir-induced formation of GPCR - Gs protein - β-arrestin megacomplexes. **a,** Atazanavir induces sustained G protein signaling in β_2_AR, M_3_R, α_1_AR and D_1_R-2mut in cAMP GloSensor assay. Data are given as Mean ± S.E.M. of 6-10 independent replicates. **b-c,** Atazanavir induces the formation of megacomplexes in β_2_AR, M_3_R, α_1_AR, D_1_R-2mut (**b**) and μOR (**c**) in NanoBiT-based miniGs-β-arrestin co-localization assay. Data are given as Mean ± S.E.M. of 8-9 independent samples. **d-e,** Both membrane-permeable antagonists (ICI-118551/metoprolol) and membrane-impermeable antagonists (CGP-12177/sotalol) can suppress isoprenaline-induced Gs signaling of β_2_AR (**d**) and β_1_AR (**e**). Data are given as Mean ± S.E.M. of 6 independent replicates.

**Extended Data Table 1.** Cryo-EM data collection, refinement and validation statistics.

| | GPR119-Gs-AR231453<br>(EMDB-62871)<br>(PDB 9L79) | GPR119-Gs-atazanavir<br>(EMDB-62880)<br>(PDB 9L80) | $\beta_1$AR-Gs-epinephrine-atazanavir<br>(EMDB-62884)<br>(PDB 9L84) |
| --- | --- | --- | --- |
| **Data collection and processing** |  |  |  |
| Magnification | 96,000 | 96,000 | 64,000 |
| Voltage (kV) | 300 | 300 | 300 |
| Electron exposure (e-/Å<sup>2</sup>) | 50 | 50 | 50 |
| Defocus range (μm) | -1.3~-1.6 | -1.3~-1.6 | -1.3~-1.6 |
| Pixel size (Å) | 0.86 | 0.86 | 1.08 |
| Symmetry imposed | C1 | C1 | C1 |
| Initial particle images (no.) | 2,817,323 | 3,109,923 | 2,255,045 |
| Final particle images (no.) | 322,758 | 106,271 | 132,798 |
| Map resolution (Å) | 2.98 | 3.33 | 3.13 |
| FSC threshold | 0.143 | 0.143 | 0.143 |
| Map resolution range (Å) | 2.55-47.75 | 2.88-18.79 | 2.75-37.22 |
| **Refinement** |  |  |  |
| Initial model used (PDB code) | 7XZ5 | 7XZ5 | 3SN6, 7BTS |
| Model resolution (Å) | 2.96 | 3.3 | 3.4 |
| FSC threshold | 0.143 | 0.143 | 0.143 |
| Model resolution range (Å) | 2.90-3.16 | 3.3-3.5 | 3.0-3.4 |
| Map sharpening *B* factor (Å<sup>2</sup>) | -143.1 | -148.3 | -114.0 |
| Model composition |  |  |  |
| Non-hydrogen atoms | 9463 | 9490 | 7472 |
| Protein residues | 1243 | 1236 | 1026 |
| Ligands | 1 | 1 | 2 |
| *B* factors (Å<sup>2</sup>) |  |  |  |
| Protein | 61.80 | 92.72 | 69.01 |
| Ligand | 92.26 | 80.90 | 102.75 |
| R.m.s. deviations |  |  |  |
| Bond lengths (Å) | 0.002 | 0.002 | 0.003 |
| Bond angles (°) | 0.547 | 0.548 | 0.671 |
| Validation |  |  |  |
| MolProbity score | 1.53 | 1.36 | 1.96 |
| Clashscore | 3.17 | 3.75 | 12.28 |
| Poor rotamers (%) | 0 | 0.41 | 0.56 |
| Ramachandran plot |  |  |  |
| Favored (%) | 93.54 | 96.79 | 94.80 |
| Allowed (%) | 6.46 | 3.21 | 5.20 |
| Disallowed (%) | 0 | 0 | 0 |

